# A contribution of novel CNVs to schizophrenia from a genome-wide study of 41,321 subjects: CNV Analysis Group and the Schizophrenia Working Group of the Psychiatric Genomics Consortium

**DOI:** 10.1101/040493

**Authors:** Christian R. Marshall, Daniel P. Howrigan, Daniele Merico, Bhooma Thiruvahindrapuram, Wenting Wu, Douglas S. Greer, Danny Antaki, Aniket Shetty, Peter A. Holmans, Dalila Pinto, Madhusudan Gujral, William M. Brandler, Dheeraj Malhotra, Zhouzhi Wang, Karin V. Fuentes Fajarado, Stephan Ripke, Ingrid Agartz, Esben Agerbo, Margot Albus, Madeline Alexander, Farooq Amin, Joshua Atkins, Silviu A. Bacanu, Richard A. Belliveau, Sarah E. Bergen, Marcelo Bertalan, Elizabeth Bevilacqua, Tim B. Bigdeli, Donald W. Black, Richard Bruggeman, Nancy G. Buccola, Randy L. Buckner, Brendan Bulik-Sullivan, William Byerley, Wiepke Cahn, Guiqing Cai, Murray J. Cairns, Dominique Campion, Rita M. Cantor, Vaughan J. Carr, Noa Carrera, Stanley V. Catts, Kimberley D. Chambert, Wei Cheng, C. Robert Cloninger, David Cohen, Paul Cormican, Nick Craddock, Benedicto Crespo-Facorro, James J. Crowley, David Curtis, Michael Davidson, Kenneth L Davis, Franziska Degenhardt, Jurgen Del Favero, Lynn E. DeLisi, Ditte Demontis, Dimitris Dikeos, Timothy Dinan, Srdjan Djurovic, Gary Donohoe, Elodie Drapeau, Jubao Duan, Frank Dudbridge, Peter Eichhammer, Johan Eriksson, Valentina Escott-Price, Laurent Essioux, Ayman H. Fanous, Kai-How Farh, Martilias S. Farrell, Josef Frank, Lude Franke, Robert Freedman, Nelson B. Freimer, Joseph I. Friedman, Andreas J. Forstner, Menachem Fromer, Giulio Genovese, Lyudmila Georgieva, Elliot S. Gershon, Ina Giegling, Paola Giusti-Rodríguez, Stephanie Godard, Jacqueline I. Goldstein, Jacob Gratten, Lieuwe de Haan, Marian L. Hamshere, Mark Hansen, Thomas Hansen, Vahram Haroutunian, Annette M. Hartmann, Frans A. Henskens, Stefan Herms, Joel N. Hirschhorn, Per Hoffmann, Andrea Hofman, Mads V. Hollegaard, David M. Hougaard, Hailiang Huang, Masashi Ikeda, Inge Joa, K Kähler Anna, René S Kahn, Luba Kalaydjieva, Juha Karjalainen, David Kavanagh, Matthew C. Keller, Brian J. Kelly, James L. Kennedy, Yunjung Kim, James A. Knowles, Bettina Konte, Claudine Laurent, Phil Lee, S. Hong Lee, Sophie E. Legge, Bernard Lerer, Deborah L. Levy, Kung-Yee Liang, Jeffrey Lieberman, Jouko Lönnqvist, Carmel M. Loughland, Patrik K.E. Magnusson, Brion S. Maher, Wolfgang Maier, Jacques Mallet, Manuel Mattheisen, Morten Mattingsdal, Robert W McCarley, Colm McDonald, Andrew M. McIntosh, Sandra Meier, Carin J. Meijer, Ingrid Melle, Raquelle I. Mesholam-Gately, Andres Metspalu, Patricia T. Michie, Lili Milani, Vihra Milanova, Younes Mokrab, Derek W. Morris, Ole Mors, Bertram Müller-Myhsok, Kieran C. Murphy, Robin M. Murray, Inez Myin-Germeys, Igor Nenadic, Deborah A. Nertney, Gerald Nestadt, Kristin K. Nicodemus, Laura Nisenbaum, Annelie Nordin, Eadbhard O’ Callaghan, Colm O’ Dushlaine, Sang-Yun Oh, Ann Olincy, Line Olsen, F. Anthony O’ Neill, Jim Van Os, Christos Pantelis, George N. Papadimitriou, Elena Parkhomenko, Michele T. Pato, Tiina Paunio, Psychosis Endophenotypes International Consortium, Diana O. Perkins, Tune H. Pers, Olli Pietiläinen, Jonathan Pimm, Andrew J. Pocklington, John Powell, Alkes Price, Ann E. Pulver, Shaun M. Purcell, Digby Quested, Henrik B. Rasmussen, Abraham Reichenberg, Mark A. Reimers, Alexander L. Richards, Joshua L. Roffman, Panos Roussos, Douglas M. Ruderfer, Veikko Salomaa, Alan R. Sanders, Adam Savitz, Ulrich Schall, Thomas G. Schulze, Sibylle G. Schwab, Edward M. Scolnick, Rodney J. Scott, Larry J. Seidman, Jianxin Shi, Jeremy M. Silverman, Jordan W. Smoller, Erik Söderman, Chris C.A. Spencer, Eli A. Stahl, Eric Strengman, Jana Strohmaier, T. Scott Stroup, Jaana Suvisaari, Dragan M. Svrakic, Jin P. Szatkiewicz, Srinivas Thirumalai, Paul A. Tooney, Juha Veijola, Peter M. Visscher, John Waddington, Dermot Walsh, Bradley T. Webb, Mark Weiser, Dieter B. Wildenauer, Nigel M. Williams, Stephanie Williams, Stephanie H. Witt, Aaron R. Wolen, Brandon K. Wormley, Naomi R Wray, Jing Qin Wu, Clement C. Zai, Wellcome Trust Case-Control Consortium, Rolf Adolfsson, Ole A. Andreassen, Douglas H.R. Blackwood, Anders D. Børglum, Elvira Bramon, Joseph D. Buxbaum, Sven Cichon, David A. Collier, Aiden Corvin, Mark J. Daly, Ariel Darvasi, Enrico Domenici, Tõnu Esko, Pablo V. Gejman, Michael Gill, Hugh Gurling, Christina M. Hultman, Nakao Iwata, Assen V. Jablensky, Erik G Jönsson, Kenneth S Kendler, George Kirov, Jo Knight, Douglas F. Levinson, Qingqin S Li, Steven A McCarroll, Andrew McQuillin, Jennifer L. Moran, Preben B. Mortensen, Bryan J. Mowry, Markus M. Nöthen, Roel A. Ophoff, Michael J. Owen, Aarno Palotie, Carlos N. Pato, Tracey L. Petryshen, Danielle Posthuma, Marcella Rietschel, Brien P. Riley, Dan Rujescu, Pamela Sklar, David St. Clair, James T.R. Walters, Thomas Werge, Patrick F. Sullivan, Michael C O’Donovan, Stephen W. Scherer, Benjamin M. Neale, Jonathan Sebat

**Affiliations:** The Centre for Applied Genomics and Program in Genetics and Genome Biology, The Hospital for Sick Children,Toronto, ON, Canada; Analytic and Translational Genetics Unit, Massachusetts General Hospital, Boston, Massachusetts 02114, USA; Stanley Center for Psychiatric Research, Broad Institute of MIT and Harvard, Cambridge, Massachusetts 02142, USA; Beyster Center for Psychiatric Genomics, University of California, San Diego, La Jolla, CA 92093, USA; Department of Psychiatry, University of California, San Diego, La Jolla, CA 92093, USA; MRC Centre for Neuropsychiatric Genetics and Genomics, Institute of Psychological Medicine and clinical Neurosciences, School of Medicine, Cardiff University, Cardiff, CF24 4HQ, UK; National Centre for Mental Health, Cardiff University, Cardiff, CF24 4HQ, UK; Department of Psychiatry, Icahn School of Medicine at Mount Sinai, New York, New York 10029, USA; Department of Genetics and Genomic Sciences, Seaver Autism Center, The Mindich Child Health & Development Institute, Icahn School of Medicine at Mount Sinai, New York, New York 10029, USA; Neuroscience Discovery and Translational Area, Pharma Research & Early Development, F. Hoffmann-La Roche Ltd, CH-4070 Basel, Switzerland; NORMENT, KG Jebsen Centre for Psychosis Research, Institute of Clinical Medicine, University of Oslo, 0424 Oslo, Norway; Department of Psychiatry, Diakonhjemmet Hospital, 0319 Oslo, Norway; Department of Clinical Neuroscience, Psychiatry Section, Karolinska Institutet, SE-17176 Stockholm, Sweden; National Centre for Register-based Research, Aarhus University, DK-8210 Aarhus, Denmark; Centre for Integrative Register-based Research, CIRRAU, Aarhus University, DK-8210 Aarhus, Denmark; The Lundbeck Foundation Initiative for Integrative Psychiatric Research, iPSYCH, Denmark; State Mental Hospital, 85540 Haar, Germany; Department of Psychiatry and Behavioral Sciences, Stanford University, Stanford, California 94305, USA; Department of Psychiatry and Behavioral Sciences, Emory University, Atlanta, Georgia 30322, USA; Department of Psychiatry and Behavioral Sciences, Atlanta Veterans Affairs Medical Center, Atlanta, Georgia 30033, USA; School of Biomedical Sciences and Pharmacy, University of Newcastle, Callaghan NSW 2308, Australia; Hunter Medical Research Institute, New Lambton, New South Wales, Australia; Virginia Institute for Psychiatric and Behavioral Genetics, Department of Psychiatry, Virginia Commonwealth University, Richmond, Virginia 23298, USA; Department of Medical Epidemiology and Biostatistics, Karolinska Institutet, Stockholm SE-17177, Sweden; Institute of Biological Psychiatry, Mental Health Centre Sct. Hans, Mental Health Services Copenhagen, DK-4000, Denmark; Department of Psychiatry, University of Iowa Carver College of Medicine, Iowa City, Iowa 52242, USA; University Medical Center Groningen, Department of Psychiatry, University of Groningen, NL-9700 RB, The Netherlands; School of Nursing, Louisiana State University Health Sciences Center, New Orleans, Louisiana 70112, USA; Center for Brain Science, Harvard University, Cambridge, Massachusetts 02138, USA; Department of Psychiatry, Massachusetts General Hospital, Boston, Massachusetts 02114, USA; Athinoula A. Martinos Center, Massachusetts General Hospital, Boston, Massachusetts 02129, USA; Department of Psychiatry, University of California at San Francisco, San Francisco, California, 94143 USA; University Medical Center Utrecht, Department of Psychiatry, Rudolf Magnus Institute of Neuroscience, 3584 Utrecht, The Netherlands; Department of Human Genetics, Icahn School of Medicine at Mount Sinai, New York, New York 10029, USA; Schizophrenia Research Institute, Sydney NSW 2010, Australia; Priority Centre for Translational Neuroscience and Mental Health, University of Newcastle, Newcastle NSW 2300, Australia; Centre Hospitalier du Rouvray and INSERM U1079 Faculty of Medicine, 76301 Rouen, France; Department of Human Genetics, David Geffen School of Medicine, University of California, Los Angeles, California 90095, USA; School of Psychiatry, University of New South Wales, Sydney NSW 2031, Australia; Royal Brisbane and Women’s Hospital, University of Queensland, Brisbane QLD 4072, Australia; Department of Computer Science, University of North Carolina, Chapel Hill, North Carolina 27514, USA; Department of Psychiatry, Washington University, St. Louis, Missouri 63110, USA; Department of Child and Adolescent Psychiatry, Assistance Publique Hospitaux de Paris, Pierre and Marie Curie Faculty of Medicine and Institute for Intelligent Systems and Robotics, Paris, 75013, France; Neuropsychiatric Genetics Research Group, Department of Psychiatry, Trinity College Dublin, Dublin 8, Ireland; University Hospital Marqués de Valdecilla, Instituto de Formación e Investigación Marqués de Valdecilla, University of Cantabria, E-39008 Santander, Spain; Centro Investigación Biomédica en Red Salud Mental, Madrid, Spain; Department of Genetics, University of North Carolina, Chapel Hill, North Carolina 27599-7264, USA; Department of Psychological Medicine, Queen Mary University of London, London E1 1BB, UK; Molecular Psychiatry Laboratory, Division of Psychiatry, University College London, London WC1E 6JJ, UK; Sheba Medical Center, Tel Hashomer 52621, Israel; Institute of Human Genetics, University of Bonn, D-53127 Bonn, Germany; Department of Genomics, Life and Brain Center, D-53127 Bonn, Germany; Applied Molecular Genomics Unit, VIB Department of Molecular Genetics, University of Antwerp, B-2610 Antwerp, Belgium; VA Boston Health Care System, Brockton, Massachusetts 02301, USA; Department of Psychiatry, Harvard Medical School, Boston, Massachusetts 02115, USA; Department of Biomedicine, Aarhus University, DK-8000 Aarhus C, Denmark; Centre for Integrative Sequencing, iSEQ, Aarhus University, DK-8000 Aarhus C, Denmark; First Department of Psychiatry, University of Athens Medical School, Athens 11528, Greece; Department of Psychiatry, University College Cork, Co. Cork, Ireland; Department of Medical Genetics, Oslo University Hospital, 0424 Oslo, Norway; Cognitive Genetics and Therapy Group, School of Psychology and Discipline of Biochemistry, National University of Ireland Galway, Co. Galway, Ireland; Department of Psychiatry and Behavioral Sciences, NorthShore University HealthSystem, Evanston, Illinois 60201, USA; Department of Psychiatry and Behavioral Neuroscience, University of Chicago, Chicago, Illinois 60637, USA; Department of Non-Communicable Disease Epidemiology, London School of Hygiene and Tropical Medicine, London WC1E 7HT, UK; Department of Psychiatry, University of Regensburg, 93053 Regensburg, Germany; Folkhälsan Research Center, Helsinki, Finland, Biomedicum Helsinki 1, Haartmaninkatu 8, FI-00290, Helsinki, Finland; National Institute for Health and Welfare, P.O. BOX 30, FI-00271 Helsinki, Finland; Department of General Practice, Helsinki University Central Hospital, University of Helsinki P.O. BOX 20, Tukholmankatu 8 B, FI-00014, Helsinki, Finland; Translational Technologies and Bioinformatics, Pharma Research and Early Development, F.Hoffman-La Roche, CH-4070 Basel, Switzerland; Mental Health Service Line, Washington VA Medical Center, Washington DC 20422, USA; Department of Psychiatry, Georgetown University, Washington DC 20057, USA; Department of Psychiatry, Virginia Commonwealth University, Richmond, Virginia 23298, USA; Department of Psychiatry, Keck School of Medicine at University of Southern California, Los Angeles, California 90033, USA; Department of Genetic Epidemiology in Psychiatry, Central Institute of Mental Health, Medical Faculty Mannheim, University of Heidelberg, Heidelberg, D-68159 Mannheim, Germany; Department of Genetics, University of Groningen, University Medical Centre Groningen, 9700 RB Groningen, The Netherlands; Department of Psychiatry, University of Colorado Denver, Aurora, Colorado 80045, USA; Center for Neurobehavioral Genetics, Semel Institute for Neuroscience and Human Behavior, University of California, Los Angeles, California 90095, USA; Division of Psychiatric Genomics, Department of Psychiatry, Icahn School of Medicine at Mount Sinai, New York, New York 10029, USA; Psychiatric and Neurodevelopmental Genetics Unit, Massachusetts General Hospital, Boston, Massachusetts 02114, USA; Departments of Psychiatry and Human Genetics, University of Chicago, Chicago, Illinois 60637 USA; Department of Psychiatry, University of Halle, 06112 Halle, Germany; Department of Psychiatry, University of Munich, 80336, Munich, Germany; Departments of Psychiatry and Human and Molecular Genetics, INSERM, Institut de Myologie, Hôpital de la Pitiè-Salpêtrière, Paris, 75013, France; Medical and Population Genetics Program, Broad Institute of MIT and Harvard, Cambridge, Massachusetts 02142, USA; Queensland Brain Institute, The University of Queensland, Brisbane, QLD 4072, Australia; Academic Medical Centre University of Amsterdam, Department of Psychiatry, 1105 AZ Amsterdam, The Netherlands; Illumina, La Jolla, California, California 92122, USA; J.J. Peters VA Medical Center, Bronx, New York, New York 10468, USA; Friedman Brain Institute, Icahn School of Medicine at Mount Sinai, New York, New York 10029, USA; School of Electrical Engineering and Computer Science, University of Newcastle, Newcastle NSW 2308, Australia; Division of Medical Genetics, Department of Biomedicine, University of Basel, Basel, CH-4058, Switzerland; Department of Genetics, Harvard Medical School, Boston, Massachusetts 02115, USA; Division of Endocrinology and Center for Basic and Translational Obesity Research, Boston Children’s Hospital, Boston, Massachusetts 02115, USA; Section of Neonatal Screening and Hormones, Department of Clinical Biochemistry, Immunology and Genetics, Statens Serum Institut, Copenhagen, DK-2300, Denmark; Department of Psychiatry, Fujita Health University School of Medicine, Toyoake, Aichi, 470-1192, Japan; Regional Centre for Clinical Research in Psychosis, Department of Psychiatry, Stavanger University Hospital, 4011 Stavanger, Norway; Centre for Medical Research, The University of Western Australia, Perth, WA 6009, Australia; School of Psychiatry and Clinical Neurosciences, The University of Western Australia, Perth, WA 6009, Australia; Department of Psychology, University of Colorado Boulder, Boulder, Colorado 80309, USA; Campbell Family Mental Health Research Institute, Centre for Addiction and Mental Health, Toronto, Ontario, M5T 1R8, Canada; Department of Psychiatry, University of Toronto, Toronto, Ontario, M5T 1R8, Canada; Institute of Medical Science, University of Toronto, Toronto, Ontario, M5S 1A8, Canada; Department of Psychiatry and Zilkha Neurogenetics Institute, Keck School of Medicine at University of Southern California, Los Angeles, California 90089, USA; Department of Child and Adolescent Psychiatry, Pierre and Marie Curie Faculty of Medicine, Paris 75013, France; Department of Psychiatry, Hadassah-Hebrew University Medical Center, Jerusalem 91120, Israel; Psychology Research Laboratory, McLean Hospital, Belmont, MA; Department of Biostatistics, Johns Hopkins University Bloomberg School of Public Health, Baltimore, Maryland 21205, USA; Department of Psychiatry, Columbia University, New York, New York 10032, USA; Department of Mental Health and Substance Abuse Services, National Institute for Health and Welfare, P.O. BOX 30, FI-00271 Helsinki, Finland; Department of Mental Health, Bloomberg School of Public Health, Johns Hopkins University, Baltimore, Maryland 21205, USA; Department of Psychiatry, University of Bonn, D-53127 Bonn, Germany; Centre National de la Recherche Scientifique, Laboratoire de Génétique Moléculaire de la Neurotransmission et des Processus Neurodégénératifs, Hôpital de la Pitié Salpêtrière, 75013, Paris, France; Department of Genomics Mathematics, University of Bonn, D-53127 Bonn, Germany; Research Unit, Sørlandet Hospital, 4604 Kristiansand, Norway; Department of Psychiatry, National University of Ireland Galway, Co. Galway, Ireland; Division of Psychiatry, University of Edinburgh, Edinburgh EH16 4SB, UK; Centre for Cognitive Ageing and Cognitive Epidemiology, University of Edinburgh, Edinburgh EH16 4SB, UK; Division of Mental Health and Addiction, Oslo University Hospital, 0424 Oslo, Norway; Massachusetts Mental Health Center Public Psychiatry Division of the Beth Israel Deaconess Medical Center, Boston, Massachusetts 02114, USA; Estonian Genome Center, University of Tartu, Tartu 50090, Estonia; School of Psychology, University of Newcastle, Newcastle NSW 2308, Australia; First Psychiatric Clinic, Medical University, Sofia 1431, Bulgaria; Eli Lilly and Company Limited, Erl Wood Manor, Sunninghill Road, Windlesham, Surrey, GU20 6PH UK; Department P, Aarhus University Hospital, DK-8240 Risskov, Denmark; Max Planck Institute of Psychiatry, 80336 Munich, Germany; Institute of Translational Medicine, University of Liverpool, Liverpool L69 3BX, UK127Munich; Cluster for Systems Neurology (SyNergy), 80336 Munich, Germany; Department of Psychiatry, Royal College of Surgeons in Ireland, Dublin 2, Ireland; King’s College London, London SE5 8AF, UK; Maastricht University Medical Centre, South Limburg Mental Health Research and Teaching Network, EURON, 6229 HX Maastricht, The Netherlands; Department of Psychiatry and Psychotherapy, Jena University Hospital, 07743 Jena, Germany; Queensland Centre for Mental Health Research, University of Queensland, Brisbane QLD 4076, Australia; Department of Psychiatry and Behavioral Sciences, Johns Hopkins University School of Medicine, Baltimore, Maryland 21205, USA; Department of Psychiatry, Trinity College Dublin, Dublin 2, Ireland; Eli Lilly and Company, Lilly Corporate Center, Indianapolis, 46285 Indiana, USA; Department of Clinical Sciences, Psychiatry, Umeå University, SE-901 87 Umeå, Sweden; DETECT Early Intervention Service for Psychosis, Blackrock, Co. Dublin, Ireland; Lawrence Berkeley National Laboratory, University of California at Berkeley, Berkeley, California 94720, USA; Centre for Public Health, Institute of Clinical Sciences, Queen’s University Belfast, Belfast BT12 6AB, UK; Institute of Psychiatry, King’s College London, London SE5 8AF, UK; Melbourne Neuropsychiatry Centre, University of Melbourne & Melbourne Health, Melbourne VIC 3053, Australia; Public Health Genomics Unit, National Institute for Health and Welfare, P.O. BOX 30, FI-00271 Helsinki, Finland; Department of Psychiatry, University of North Carolina, Chapel Hill, North Carolina 27599-7160, USA; Center for Biological Sequence Analysis, Department of Systems Biology, Technical University of Denmark, DK-2800, Denmark; Institute for Molecular Medicine Finland, FIMM, University of Helsinki, P.O. BOX 20 FI-00014, Helsinki, Finland; Department of Epidemiology, Harvard School of Public Health, Boston, Massachusetts 02115, USA; Department of Psychiatry, University of Oxford, Oxford, OX3 7JX, UK; Institute for Multiscale Biology, Icahn School of Medicine at Mount Sinai, New York, New York 10029, USA; Neuroscience Therapeutic Area, Janssen Research and Development, Raritan, New Jersey 08869, USA; Department of Psychiatry and Psychotherapy, University of Göttingen, 37073 Göttingen, Germany; Psychiatry and Psychotherapy Clinic, University of Erlangen, 91054 Erlangen, Germany; Hunter New England Health Service, Newcastle NSW 2308, Australia; Division of Cancer Epidemiology and Genetics, National Cancer Institute, Bethesda, Maryland 20892, USA; Research and Development, Bronx Veterans Affairs Medical Center, New York, New York 10468, USA; Wellcome Trust Centre for Human Genetics, Oxford, OX3 7BN, UK; Department of Medical Genetics, University Medical Centre Utrecht, Universiteitsweg 100, 3584 CG, Utrecht, The Netherlands; Berkshire Healthcare NHS Foundation Trust, Bracknell RG12 1BQ, UK; Department of Psychiatry, University of Oulu, P.O. BOX 5000, 90014, Finland; University Hospital of Oulu, P.O. BOX 20, 90029 OYS, Finland; Molecular and Cellular Therapeutics, Royal College of Surgeons in Ireland, Dublin 2, Ireland; Health Research Board, Dublin 2, Ireland; University College London, London WC1E 6BT, UK; Department of Neuroscience, Icahn School of Medicine at Mount Sinai, New York, New York 10029, USA; Institute of Neuroscience and Medicine (INM-1), Research Center Juelich, 52428 Juelich, Germany; Social, Genetic and Developmental Psychiatry Centre, Institute of Psychiatry, King’s College London, London, SE5 8AF, UK; Department of Genetics, The Hebrew University of Jerusalem, 91905 Jerusalem, Israel; The Perkins Institute for Medical Research, The University of Western Australia, Perth, WA 6009, Australia; Centre for Clinical Research in Neuropsychiatry, School of Psychiatry and Clinical Neurosciences, The University of Western Australia, Medical Research Foundation Building, Perth WA 6000, Australia; Center for Human Genetic Research and Department of Psychiatry, Massachusetts General Hospital, Boston, Massachusetts 02114, USA; Department of Functional Genomics, Center for Neurogenomics and Cognitive Research, Neuroscience Campus Amsterdam, VU University, Amsterdam 1081, The Netherlands; Department of Complex Trait Genetics, Neuroscience Campus Amsterdam, VU University Medical Center Amsterdam, Amsterdam 1081, The Netherlands; Department of Child and Adolescent Psychiatry, Erasmus University Medical Centre, Rotterdam 3000, The Netherlands; University of Aberdeen, Institute of Medical Sciences, Aberdeen, AB25 2ZD, UK; Department of Clinical Medicine, University of Copenhagen, Copenhagen 2200, Denmark; Department of Molecular Genetics and McLaughlin Centre, University of Toronto, Toronto, Ontario, Canada; Department of Cellular and Molecular Medicine, University of California, San Diego, La Jolla, CA 92093, USA

## Abstract

Genomic copy number variants (CNVs) have been strongly implicated in the etiology of schizophrenia (SCZ). However, apart from a small number of risk variants, elucidation of the CNV contribution to risk has been difficult due to the rarity of risk alleles, all occurring in less than 1% of cases. We sought to address this obstacle through a collaborative effort in which we applied a centralized analysis pipeline to a SCZ cohort of 21,094 cases and 20,227 controls. We observed a global enrichment of CNV burden in cases (OR=1.11, P=5.7e^−15^), which persisted after excluding loci implicated in previous studies (OR=1.07, P=1.7e^−6^). CNV burden is also enriched for genes associated with synaptic function (OR = 1.68, P = 2.8e^−11^) and neurobehavioral phenotypes in mouse (OR = 1.18, P=7.3e^−5^). We identified genome-wide significant support for eight loci, including 1q21.1, 2p16.3 (NRXN1), 3q29, 7q11.2, 15q13.3, distal 16p11.2, proximal 16p11.2 and 22q11.2. We find support at a suggestive level for nine additional candidate susceptibility and protective loci, which consist predominantly of CNVs mediated by non-allelic homologous recombination (NAHR).

## Introduction

Studies of genomic copy number variation (CNV) have established a role for rare genetic variants in the etiology of SCZ ^1^. There are three lines of evidence that CNVs contribute to risk for SCZ: genome-wide enrichment of rare deletions and duplications in SCZ cases relative to controls ^2,3^, a higher rate of *de novo* CNVs in cases relative to controls^4-6^, and association evidence implicating a small number of specific loci (**Extended data table 1**). All CNVs that have been implicated in SCZ are rare in the population, but confer significant risk (odds ratios 2-60).

To date, CNVs associated with SCZ have largely emerged from mergers of summary data for specific candidate loci ^7-9^; yet even the largest genome-wide scans (sample sizes typically <10,000) remain under-powered to robustly confirm genetic association for the majority of pathogenic CNVs reported so far, particularly for those with low frequencies (<0.5% in cases) or intermediate effect sizes (odds ratios 2-10). It is important to address the low power of systematic CNV studies with larger samples given that this type of mutation has already proven useful for highlighting some aspects of SCZ related biology ^6,10-13^.

The limited statistical power provided by small samples is a significant obstacle in studies of rare and common genetic variation. In response, global collaborations have been formed in order to attain large sample sizes, as exemplified by the study of the Schizophrenia Working Group of the Psychiatric Genomics Consortium (PGC) in which 108 independent schizophrenia associated loci were identified ^14^. Recognizing the need for similarly large samples in studies of CNVs for psychiatric disorders, we formed the PGC CNV Analysis Group. Our goal was to enable large-scale analyses of CNVs in psychiatry using centralized and uniform methodologies for CNV calling, quality control, and statistical analysis. Here, we report the largest genome-wide analysis of CNVs for any psychiatric disorder to date, using datasets assembled by the Schizophrenia Working Group of the PGC.

## Data processing and meta-analytic methods

Raw intensity data were obtained from 57,577 subjects from 43 separate datasets (**Extended data table 2**). After CNV calling and quality control (QC), 41,321 subjects were retained for analysis. In large datasets derived from multiple studies, variability in CNV detection between studies and array platforms presents a significant challenge. To minimize the technical variability across different studies, we developed a centralized pipeline for systematic calling of CNVs for Affymetrix and Illumina platforms. (**Methods** and **Extended data figure 1**). The pipeline included multiple CNV callers run in parallel. Data from Illumina platforms were processed using PennCNV ^15^ and iPattern ^16^. Data from Affymetrix platforms were analyzed using PennCNV and Birdsuite ^17^.Two additional methods, iPattern and C-score ^18^, were applied to data from the Affymetrix 6.0 platform. The CNV calls from each program were converted to a standardized format and a consensus call set was constructed by merging CNV outputs at the sample level. Only CNV segments that were detected by all algorithms were retained. We performed rigorous QC at the platform level to exclude samples with poor probe intensity and/or an excessive CNV load (number and length). Larger CNVs that appeared to be fragmented were merged and retained. CNVs spanning centromeres or those with >50% overlap with segmental duplications or regions prone to VDJ recombination (e.g., immunoglobulin or T cell receptor loci) were excluded. A final set of rare, high quality CNVs was defined as those >20kb in length, at least 10 probes, and <1% MAF.

Genetic associations were investigated by case-control tests of CNV burden at four levels: (1) genome-wide (2) pathways, (3) genes, and (4) probes. Analyses controlled for SNP-derived principal components, sex, genotyping platform, and individual-level probe intensity. Multiple-testing thresholds for genome-wide significance were estimated from family-wise error rates drawn from permutation

## Genome wide analysis of CNV burden reveals an enrichment of ultra-rare variants

An elevated burden of rare CNVs has been well established among SCZ cases^2^. We applied our meta-analytic framework to measure the consistency of overall CNV burden across the genotyping platforms, and whether a measurable amount of CNV burden persists outside of previously implicated CNV regions. Consistent with previous estimates, the overall CNV burden is significantly greater among SCZ cases when measured as total Kb covered (OR=1.12, p = 5.7e^−15^), genes affected (OR=1.21, p = 6.6e^−21^), or CNV number (OR=1.03, p = 1e^−3^). Focusing on genes affected by CNV, our strongest signal of enrichment, the effect size is consistent across all genotyping platforms (**Figure 1A**). When we split by CNV type, the effect size for copy number losses (OR=1.40, p = 4e^−16^) is greater than for gains (OR=1.12, p = 2e^−7^) (**Extended data figures 2**-**3**). Partitioning by CNV frequency (based on 50% reciprocal overlap with the full call set, **Methods**), CNV burden is enriched among cases across a range of frequencies, up to counts of 80 (MAF = 0.2%) in the combined sample (**Figure 1B**).

**Figure 1.**
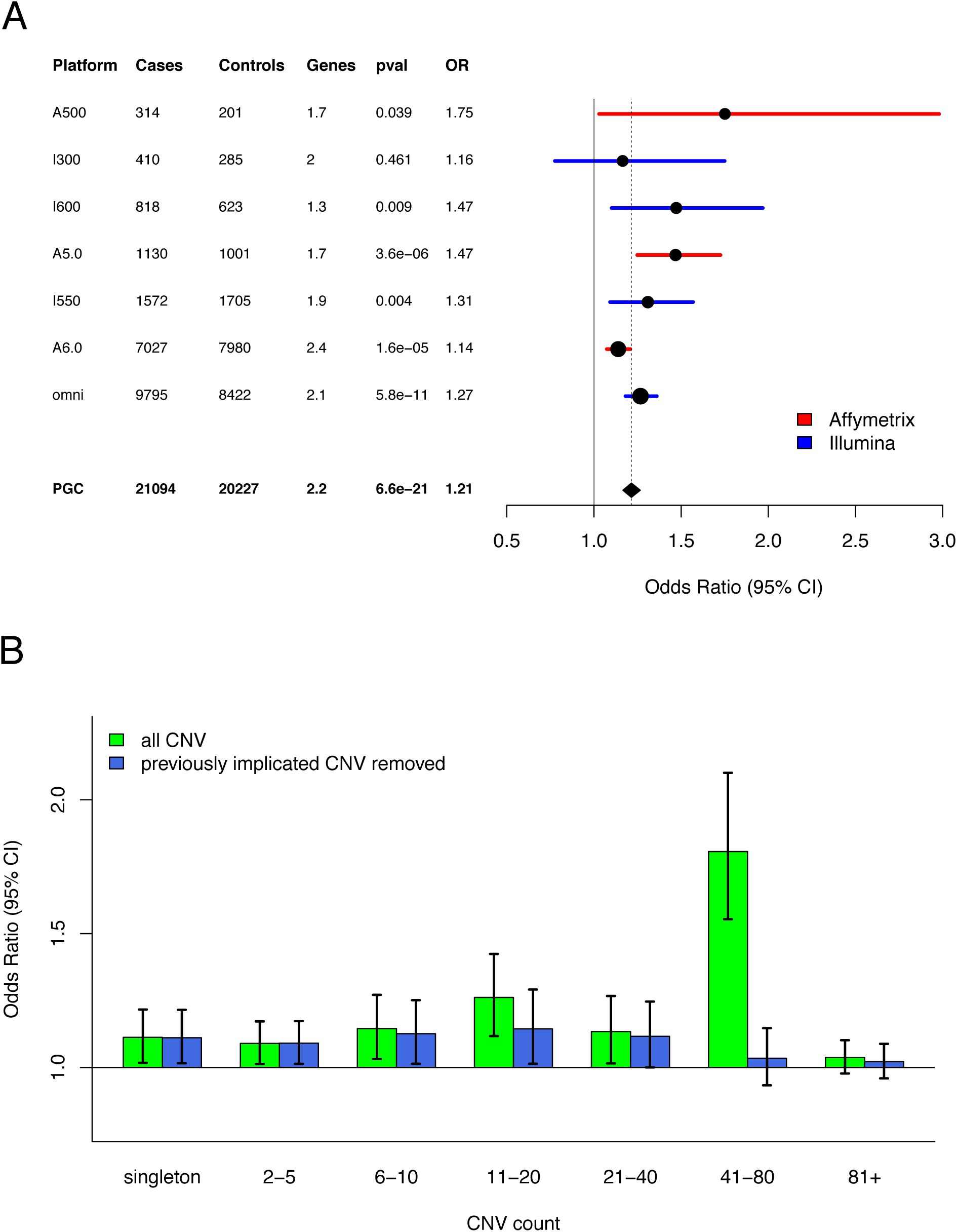
CNV Burden. **(A)** Forest plot of CNV burden (measured here as genes affected by CNV), partitioned by genotyping platform, with the full PGC sample at the bottom. CNV burden is calculated by combining CNV gains and losses. Case and control counts are listed, and “genes” is the rate of genes affected by CNV in controls. Burden tests use a logistic regression model predicting SCZ case/control status by CNV burden along with covariates (see methods). The odds ratio is the exponential of the logistic regression coefficient, and odds ratios above one predict increased SCZ risk. **(B)** CNV burden partitioned by CNV frequency. For reference, a CNV with MAF 0.1% in the PGC sample would have ˜41 CNVs. Using the same model as above, each CNV was placed into a single CNV frequency category based on a 50% reciprocal overlap with other CNVs. CNV burden with inclusion of all CNVs are shown in green, whereas CNV burden excluding previously implicated CNV loci are shown in blue.

A primary question in this study is the contribution of novel loci to the excess CNV burden in cases. After removing nine previously implicated CNV loci (where reported *p*-values exceed our designated multiple testing threshold, **Extended data table 1**), excess CNV burden in SCZ remains significantly enriched (genes affected OR=1.11, p = 1.3e^−7^, **Figure 1B**). CNV burden also remained significantly enriched after removal of all reported loci from **Extended data table 1**, but the effect-size was greatly reduced (OR = 1.08) compared to the enrichment overall (OR = 1.21). When we partition CNV burden by frequency, we find that much of the previously unexplained signal is restricted to comparatively rare events (i.e., MAF < 0.1%, **Figure 1B**).

## Gene-set (pathway) burden

We assessed whether CNV burden was concentrated within defined sets of genes involved in neurodevelopment or neurological function. A total of 36 gene-sets were evaluated (for a description see **Extended data table 3**), consisting of gene-sets representing neuronal function, synaptic components and neurological and neurodevelopmental phenotypes in human (19 sets), gene-sets based on brain expression patterns (7 sets), and human orthologs of mouse genes whose disruption causes phenotypic abnormalities, including neurobehavioral and nervous system abnormality (10 sets). Some gene-sets can be considered “negative controls”, including genes not expressed in brain (1 set) or associated with abnormal phenotypes in mouse organ systems unrelated to brain (7 sets). We mapped CNVs to genes if they overlapped by at least one exonic bp.

Gene-set burden was tested using logistic regression deviance test ^6^. In addition to using the same covariates included in genome-wide burden analysis, we controlled for the total number of genes per subject spanned by rare CNVs to account for signal that merely reflects the global enrichment of CNV burden in cases ^19^. Multiple-testing correction (Benjamini-Hochberg False Discovery Rate, BH-FDR) was performed separately for each gene-set group and CNV type (gains, losses). After multiple test correction (Benjamini-Hochberg FDR ≤ 10%) 15 gene-sets were enriched for rare loss burden in cases and 4 for rare gains in cases, all of which are brain-related gene sets (**Figure 2**).

**Figure 2:**
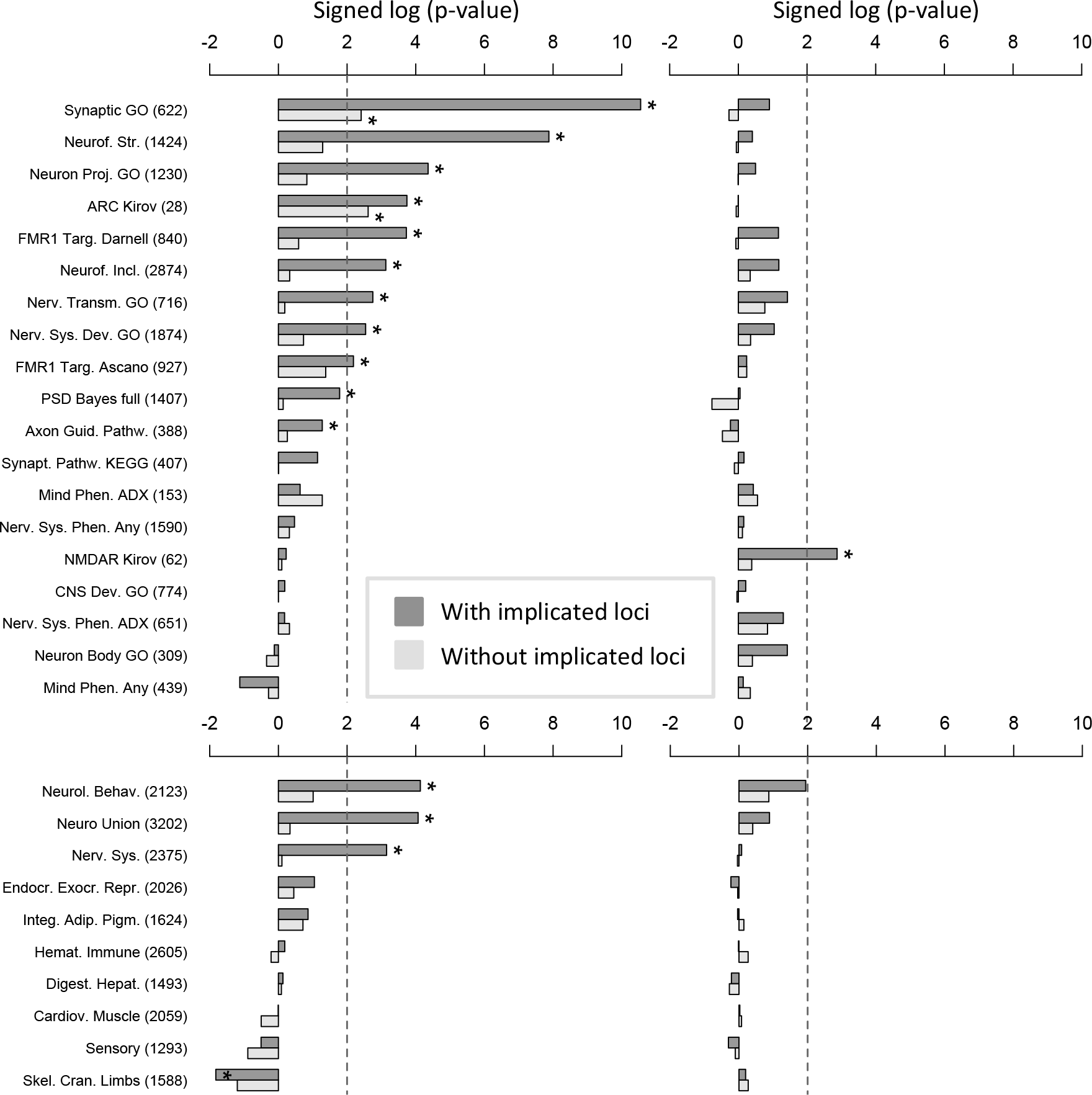
Gene-set Burden. Gene-set burden test results for rare losses (a, c) and gains (b, d); frames a-b display gene-sets for neuronal function, synaptic components, neurological and neurodevelopmental phenotypes in human; frames c-d display gene-sets for human homologs of mouse genes implicated in abnormal phenotypes (organized by organ systems); both are sorted by −log 10 of the logistic regression deviance test p-value multiplied by the beta coefficient sign, obtained for rare losses when including known loci. Gene-sets passing the 10% BH-FDR threshold are marked with “*”. Gene-sets representing brain expression patterns were omitted from the figure because only a few were significant (losses: 1, gains: 3).

Of the 15 sets significant for losses, the majority consist of synaptic or other neuronal components (9 sets) from gene-set group (a); in particular, “GO synaptic” (GO:0045202) and “ARC complex” rank first based on statistical significance and effect-size respectively (“GO synaptic” deviance test p-value = 2.8e-11, “ARC complex” regression odds-ratio > 1.8, **Figure 2a**). Losses in cases were also significantly enriched for genes involved in nervous system or behavioral phenotypes in mouse but not for gene-sets related to other organ system phenotypes (**Figure 2c**). To account for dependency between synaptic and neuronal gene-sets, we re-tested loss burden following a step-down logistic regression approach, ranking gene-sets based on significance or effect size (**Extended data table 4**). Only GO synaptic and ARC complex were significant in at least one of the two step-down analyses, suggesting that burden enrichment in the other neuronal categories is mostly accounted by the overlap with synaptic genes. Following the same approach, the mouse neurological/neurobehavioral phenotype set remained nominally significant, pointing to the existence of additional signal not captured by the synaptic set. Pathway enrichment was less pronounced for duplications, consistent with the smaller burden effects for this class of CNV. Duplication burden was significantly enriched for NMDA receptor complex, highly brain-expressed genes, medium/low brain-expressed genes and prenatally expressed brain genes (**Figure 2b**).

Given that synaptic gene sets were robustly enriched for deletions in cases, and with an appreciable contribution from loci that have not been strongly associated with SCZ previously, pathway-level interactions of these sets were further investigated. A protein-interaction network was seeded using the synaptic and ARC complex genes that were intersected by rare deletions in this study (**Figure 3**). A graph of the network highlights multiple subnetworks of synaptic proteins including pre-synaptic adhesion molecules (NRXN1, NRXN3), post-synaptic scaffolding proteins (DLG1, DLG2, DLGAP1, SHANK1, SHANK2), glutamatergic ionotropic receptors (GRID1, GRID2, GRIN1, GRIA4), and complexes such as Dystrophin and its synaptic interacting proteins (DMD, DTNB, SNTB1, UTRN). A subsequent test of the Dystrophin glycoprotein complex (DGC) revealed that deletion burden of the synaptic DGC proteins (intersection of “GO DGC” GO:0016010 and “GO synapse” GO:0045202) was enriched in cases (Deviance test P = 0.05), but deletion burden of the full DGC was not significant (P = 0.69).

**Figure 3:**
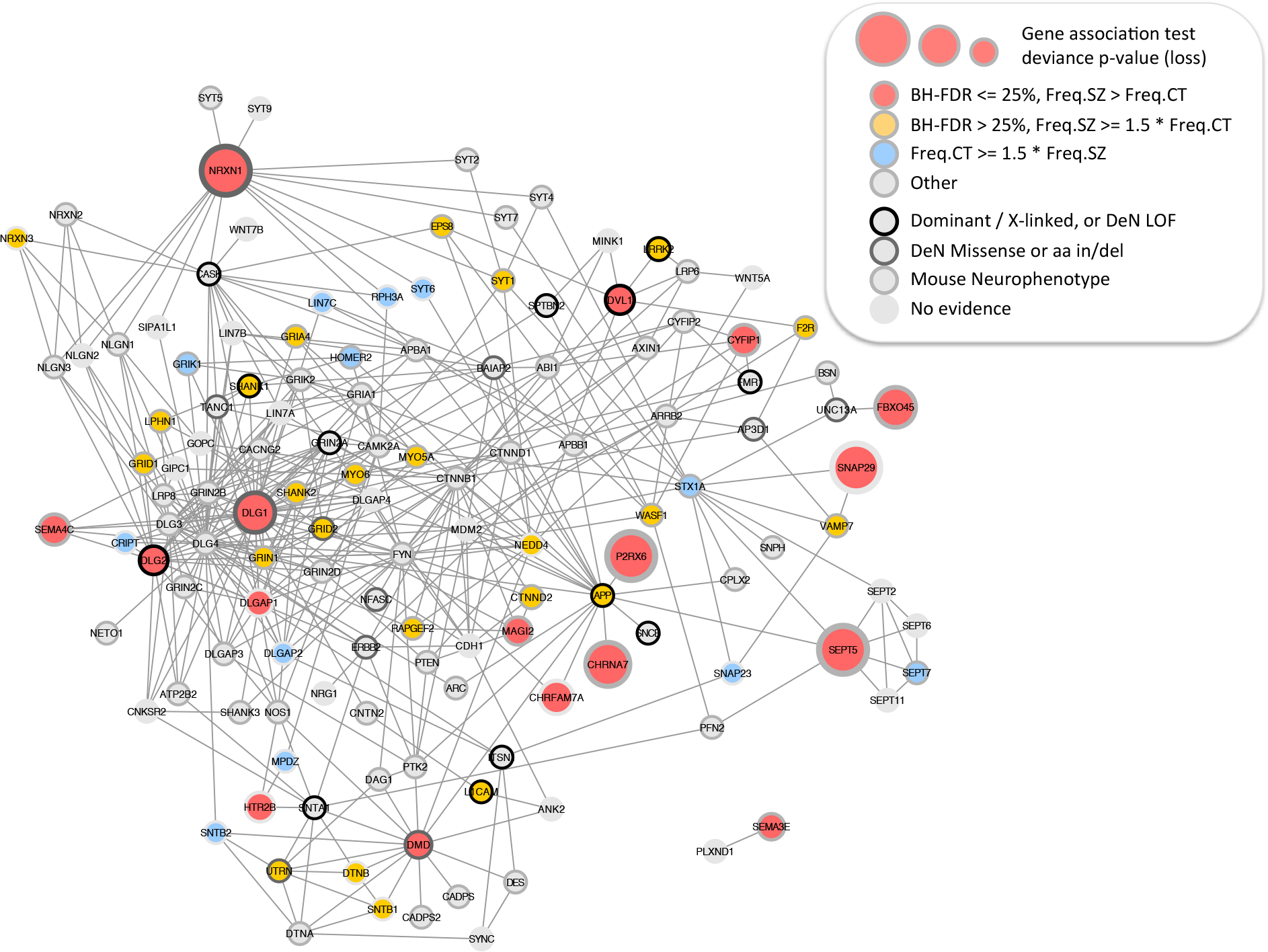
Protein Interaction Network for Synaptic Genes. Synaptic and ARC-complex genes intersected by a rare loss in at least 4 case or control subjects and with genie burden Benjamini-Hochberg FDR <= 25% (red discs) were used to query GeneMANIA^32^ and retrieve additional protein interaction neighbors, resulting in a network of 136 synaptic genes. Genes are depicted as disks; disk centers are colored based on rare loss frequency (Freq.SZ and Freq.CT) being prevalent in cases or controls; disk borders are colored to mark (i) gene implication in human dominant or X-linked neurological or neurodevelopmental phenotype, (ii) *de novo* mutation (DeN) reported by Fromer et al.^28^, split between LOF (frameshift, stop-gain, core splice site) and missense or amino acid insertion / deletion, (iii) implication in mouse neurobehavioral abnormality. Pre-synaptic adhesion molecules (NRXN1, NRXN3), post-synaptic scaffolds (DLG1, DLG2, DLGAP1, SHANK1, SHANK2) and glutamatergic ionotropic receptors (GRID1, GRID2, GRIN1, GRIA4) constitute a highly connected subnetwork with more losses in cases than controls.

## Gene CNV burden

To define specific loci that confer risk for SCZ, we tested CNV burden at the level of individual genes, using logistic regression deviance test and the same covariates included in genome-wide burden analysis. To correctly account for large CNVs that affect multiple genes, we aggregated adjacent genes into single loci if their copy number was highly correlated across subjects. CNVs were mapped to genes if they overlapped one or more exons. The criterion for genome-wide significance used the Family-Wise Error Rate (FWER) < 0.05. The criterion for suggestive evidence used a Benjamini-Hochberg False Discovery Rate (BH-FDR) < 0.05.

Of nineteen independent CNV loci with gene-based BH-FDR < 0.05, two were excluded based on CNV calling accuracy or evidence of a batch effect (**Supplementary Information**). The seventeen loci that remained after these additional QC steps are listed in **Table 1**. P-values for this summary table were obtained by re-running our statistical model across the entire region (**Supplementary Results**). These seventeen loci represent a set of novel (n=6), previously reported (n=4), and previously implicated (n=7) loci. Manhattan plots of the gene association for losses and gains are provided in **Figure 4**. A permutation-based false discovery rate yielded similar estimates to BH-FDR.

**Figure 4:**
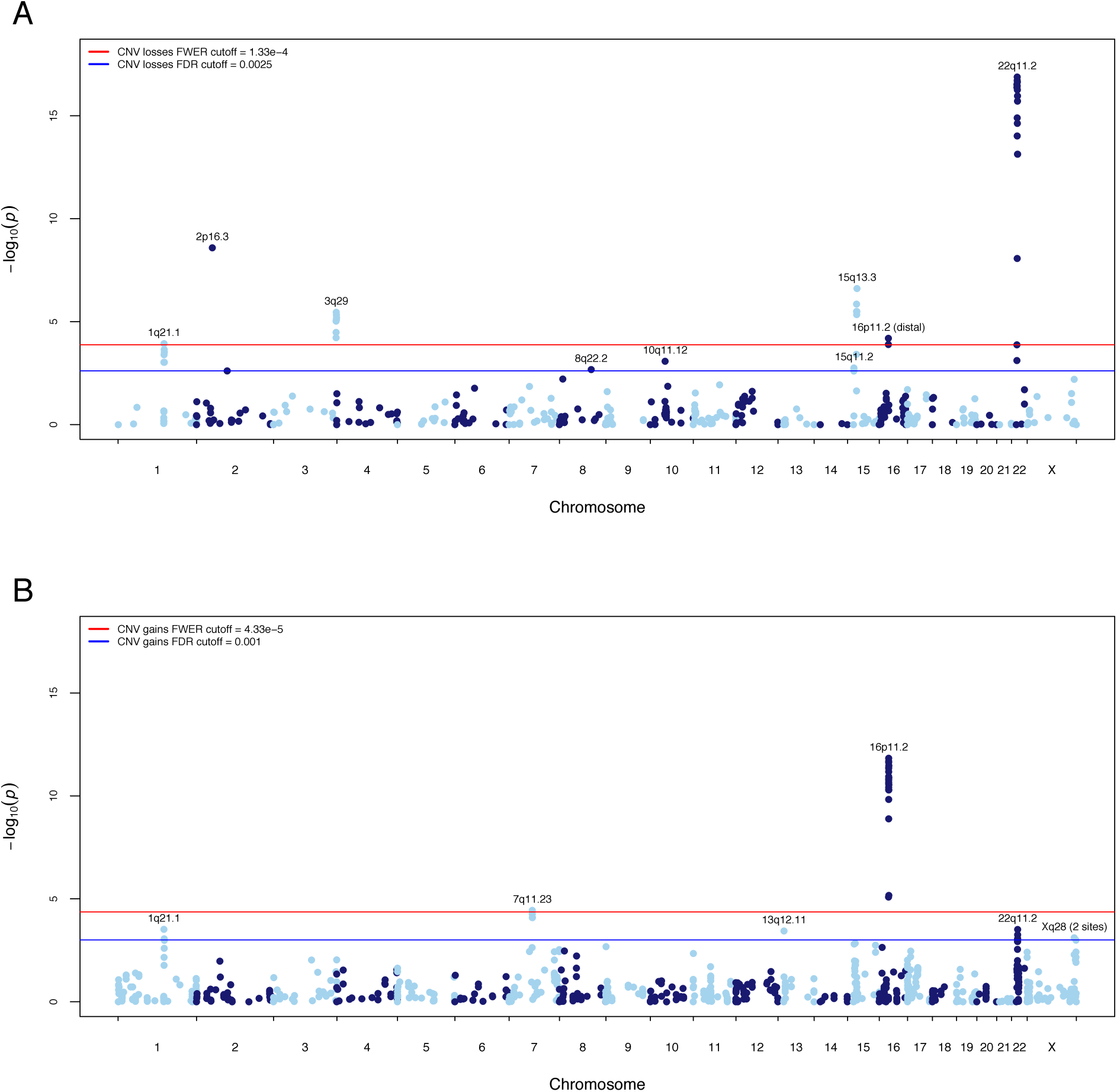
Gene Based Manhattan Plot. Manhattan plot displaying the –log10 deviance *p*-value for **(A)** CNV losses and **(B)** CNV gains the gene-based test. *P*-value cutoffs corresponding to FWER < 0.05 and BH-FDR < 0.05 are highlighted in red and blue, respectively. Loci significant after multiple test correction are labeled.

**Table 1:** Significant CNV loci from gene-based association test. All seventeen loci listed contain at least one gene with Benjamini-Hochberg false discovery rate (BH-FDR) < 0.05 in the gene-based test, with eight loci containing at least one gene surpassing the family-wise error rate (FWER) < 0.05. Genomic positions listed are using hg18 coordinates. For putative CNV mechanisms, non-allelic homologous recombination (NAHR) and non-homologous end joining (NHEJ) are listed as the likely genomic feature driving CNV formation at each locus. Regional *p*-values and odds ratios listed are from a regional test at each locus, where we combine CNV overlapping the implicated region and run the same test as used for each gene (logistic regression with covariates and deviance test *p*-value).

| CHR | BP1 | BP2 | Locus (GENE) | Putative CNV Mechanism | CNV test | Direction | FWER | BH-FDR | Cases | Controls | Regional *p*-value | Odds Ratio [95% C |
| --- | --- | --- | --- | --- | --- | --- | --- | --- | --- | --- | --- | --- |
| 22 | 17,400,000 | 19,750,000 | 22q11.21 | NAHR | loss | risk | yes | 3.54E-15 | 64 | 1 | 5.70E-18 | 67.7 [9.3-492.8] |
| 16 | 29,560,000 | 30,110,000 | 16p11.2 (proximal) | NAHR | gain | risk | yes | 5.82E-10 | 70 | 7 | 2.52E-12 | 9.4 [4.2-20.9] |
| 2 | 50,000,992 | 51,113,178 | 2p16.3 (**NRXN1**) | NHEJ | loss | risk | yes | 3.52E-07 | 35 | 3 | 4.92E-09 | 14.4 [4.2-46.9] |
| 15 | 28,920,000 | 30,270,000 | 15q13.3 | NAHR | loss | risk | yes | 2.22E-05 | 28 | 2 | 2.13E-07 | 15.6 [3.7-66.5] |
| 1 | 144,646,000 | 146,176,000 | 1q21.1 | NAHR | loss+gain | risk | yes | 0.00011 | 60 | 14 | 1.50E-06 | 3.8 [2.1-6.9] |
| 3 | 197,230,000 | 198,840,000 | 3q29 | NAHR | loss | risk | yes | 0.00024 | 16 | 0 | 1.86E-06 | NA [0-Inf] |
| 16 | 28,730,000 | 28,960,000 | 16p11.2 (distal) | NAHR | loss | risk | yes | 0.0029 | 11 | 1 | 5.52E-05 | 20.6 [2.6-162.2] |
| 7 | 72,380,000 | 73,780,000 | 7q11.23 | NAHR | gain | risk | yes | 0.0048 | 16 | 1 | 1.68E-04 | 16.1 [3.1-125.7] |
| X | 153,800,000 | 154,225,000 | Xq28 (distal) | NAHR | gain | risk | no | 0.049 | 18 | 2 | 3.61E-04 | 8.9 [2.0-39.9] |
| 22 | 17,400,000 | 19,750,000 | 22q11.21 | NAHR | gain | protective | no | 0.024 | 3 | 16 | 4.54E-04 | 0.15 [0.04-0.52] |
| 7 | 64,476,203 | 64,503,433 | 7q11.21 (**ZNF92**) | NAHR | loss+gain | protective | no | 0.033 | 131 | 180 | 6.71E-04 | 0.66 [0.52-0.84] |
| 13 | 19,309,593 | 19,335,773 | 13q12.11 (**ZMYM5**) | NHAR | gain | protective | no | 0.024 | 15 | 38 | 7.91E-04 | 0.36 [0.19-0.67] |
| X | 148,575,477 | 148,580,720 | Xq28 (**MAGEA11**) | NAHR | gain | protective | no | 0.044 | 12 | 36 | 1.06E-03 | 0.35 [0.18-0.68] |
| 15 | 20,350,000 | 20,640,000 | 15q11.2 | NAHR | loss | risk | no | 0.044 | 98 | 50 | 1.34E-03 | 1.8 [1.2-2.6] |
| 9 | 831,690 | 959,090 | 9p24.3 (**DMRT1**) | NHEJ | loss+gain | risk | no | 0.049 | 13 | 1 | 1.35E-03 | 12.4 [1.6-98.1] |
| 8 | 100,094,670 | 100,958,984 | 8q22.2 (**VPS13B**) | NHEJ | loss | risk | no | 0.048 | 7 | 1 | 1.74E-03 | 14.5 [1.7-122.2] |
| 7 | 158,145,959 | 158,664,998 | 7p36.3<br>(**VIPR2/WDR60**) | NAHR | loss+gain | risk | no | 0.046 | 20 | 6 | 5.79E-03 | 3.5 [1.3-9.0] |

Eight loci attain genome-wide significance, including copy number losses at 1q21.1, 2p16.3 (NRXN1), 3q29, 15q13.3, 16p11.2 (distal) and 22q11.2 along with gains at 7q11.23 and 16p11.2 (proximal). An additional nine loci meet criterion for suggestive association. Based on our estimation of False Discovery Rates (BH and permutations), we expect to observe less than two associations meeting suggestive criteria by chance.

## Probe level CNV burden

With our current sample size and uniform CNV calling, many individual CNV loci can be tested with adequate power at the probe level, potentially facilitating discovery at a finer grain than locus-wide tests. Tests for association were performed at each CNV breakpoint using the residuals of case-control status after controlling for analysis covariates, with significance determined through permutation. Results for losses and gains are shown in **Extended data figure 4**. Four independent CNV loci surpass genome-wide significance, all of which were also identified in the gene-based test, including the 15q13.2-13.3 and 22q11.21 deletions, 16p11.2 duplication, and 1q21.1 deletion and duplication. While these loci represent less than half of the previously implicated SCZ loci, we do find support for all loci where the association originally reported meets the criteria for genome-wide correction in this study. We examined association among all previously reported loci showing association to SCZ, including 12 CNV losses and 20 CNV gains (**Extended data table 5**), and 14 of the 33 loci were associated with SCZ at *p* < .05.

When a probe-level test is applied, associations at some loci become better delineated. For instance, The *NRXN1* gene at 2p16.3 is a CNV hotspot, and exonic deletions of this gene are significantly enriched in SCZ^9,20^. In this large sample, we observe a high density of “nonrecurrent” deletion breakpoints in cases and controls. The probe-level Manhattan plot reveals a saw tooth pattern of association, where peaks correspond to transcriptional start sites and exons of *NRXN1* (**Figure 5**). This example highlights how, with high diversity of alleles at a single locus, the association peak may become more refined, and in some cases converge toward individual functional elements. Similarly, a high density of duplication breakpoints at previously reported SCZ risk loci on 16p13.2 (http://bit.ly/1NPgIuq) and 8q11.23 (http://bit.ly/1PwdYTt) exhibit patterns of association that better delineate genes in these regions.

**Figure 5:**
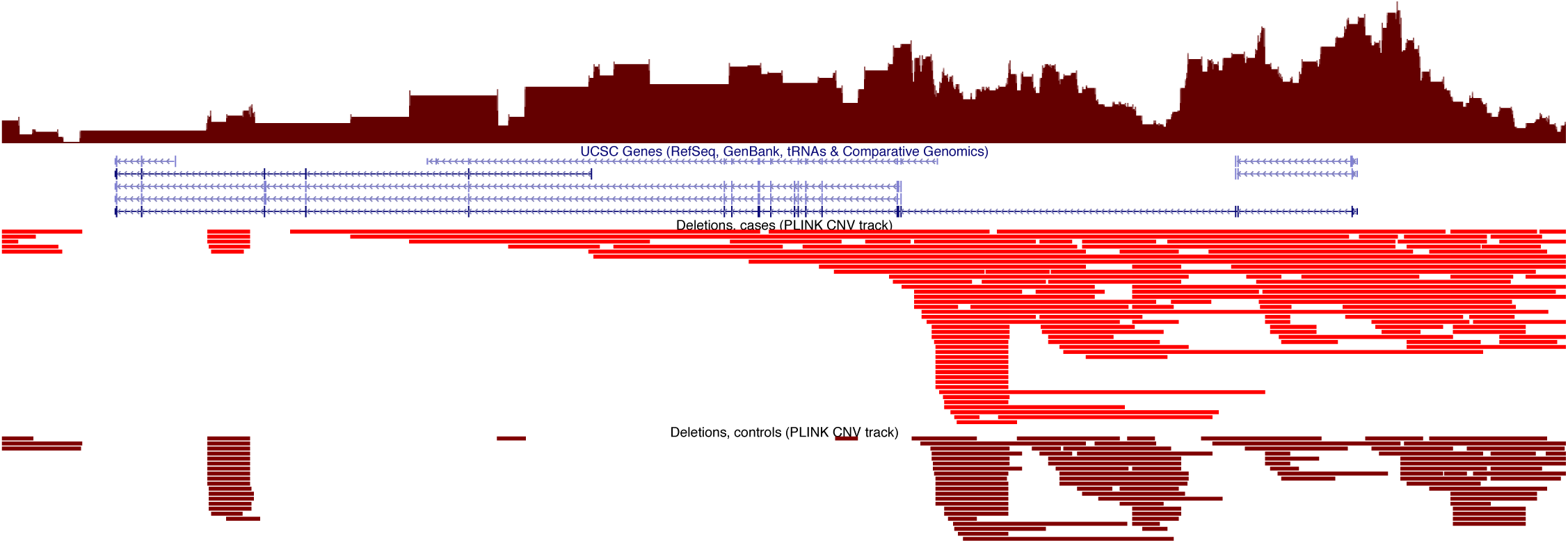
Manhattan plot of probe-level associations across the Neurexin-1 locus. Empirical *p*-values at each deletion breakpoint reveal a saw-tooth pattern of association. Predominant peaks correspond to exons and transcriptional start sites of NRXN1 isoforms.

[the above URLs link to a <u>PGC CNV browser</u> display of the respective genomic regions. The browser can also be accessed directly at the following URL http://pgc.tcag.ca/gb2/gbrowse/pgchg18/]

## Novel risk loci are predominantly NAHR-mediated CNVs

Many CNV loci that have been strongly implicated in human disease are hotspots for non-allelic homologous recombination (NAHR), a process which in most cases is mediated by flanking segmental duplications ^21^. Consistent with the importance of NAHR in generating CNV risk alleles for schizophrenia, most of the loci in **Table 1** are flanked by segmental duplications. After excluding loci that have been implicated in previous studies, we investigated whether NAHR mutational mechanisms were also enriched among novel associated CNVs. We defined a CNV as “NAHR” when both the start and end breakpoint is located within a segmental duplication. Across all loci with FDR < 0.05 in the gene-base burden test, NAHR-mediated CNVs were significantly enriched, 6.03-fold (P=0.008; **Extended data figure 5**), when compared to a null distribution determined by randomizing the genomic positions of associated genes (**Supplemental Material**). These results suggest that novel SCZ CNVs tend to occur in regions prone to high rates of recurrent mutation.

## Discussion

The present study of the PGC SCZ CNV dataset includes the majority of all microarray data that has been generated in genetic studies of SCZ to date. In this, the best body of evidence to date with which to evaluate CNV associations, we find definitive evidence for eight loci and we find significant evidence for a contribution from novel CNVs conferring both risk and protection. The complete results, including CNV calls and statistical evidence at the gene or probe level, can be viewed using the PGC CNV browser (URLs). Our data suggest that the novel risk loci that can be detected with current genotyping platforms lie at the ultra-rare end of the frequency spectrum and still larger samples will be needed to identify them at convincing levels of statistical evidence.

Collectively, the eight SCZ risk loci that surpass genome-wide significance are carried by a small fraction (1.4%) of SCZ cases in the PGC sample. We estimate 0.85% of the variance in SCZ liability is explained by carrying a CNV risk allele within these loci (**Supplementary Results**). As a comparison, 3.4% of the variance in SCZ liability is explained by the 108 genome-wide significant loci identified in the companion PGC GWAS analysis. Combined, the CNV and SNP loci that have been identified to date explain a small proportion (<5%) of heritability.

The large dataset here provides an opportunity to evaluate the strength of evidence for a variety of loci where an association with schizophrenia has been reported previously. Of 33 published findings from the recent literature, we find evidence for 14 loci (P < 0.05, **Extended data table 5**); thus, nearly half of the existing candidate loci are supported by our data. However we also find a lack of evidence for many. A lack of strong evidence in this dataset (which includes samples that overlap with many of the previous studies) may in some cases simply reflect that statistical power is limited for very rare variants, even in large samples. However, it is likely that some of these original findings represent spurious associations. Indeed, the loci that are not supported by our data consist largely of loci for which the original statistical evidence was modest (**Extended data table 5**). Thus, our results help to refine the list of promising candidate CNVs. Continued efforts to evaluate the growing number of candidate variants has considerable value for directing future research efforts focused on specific loci.

Novel candidate loci meeting suggestive criteria in this study highlight strong candidate loci that have not been previously implicated in SCZ. Two such associations are located on the X chromosome in a region of Xq28 that is highly prone to recurrent rearrangements ^22-24^ (**Extended data figure 6**). Gains at the distal Xq28 locus are enriched in cases in this study; similar duplications have been reported in association with intellectual disability, while reciprocal deletions of this region are associated with embryonic lethality in males ^25^. Duplications at the proximal Xq28 locus, including a single gene *MAGEA11*, are enriched in controls in this study, and to our knowledge have not been documented in other disorders.

We observed multiple “protective” CNVs that showed a suggestive enrichment in controls, including duplications of 22q11.2, *MAGEA11*, and *ZMYM5* along with deletions and duplications of *ZNF92*. No protective effects were significant after genome-wide correction. Moreover, a rare CNV that confers reduced risk for SCZ may not confer a general protection from neurodevelopmental disorders. For example, microduplications of 22q11.2 appear to confer protection from SCZ ^26^; however, such duplications have been shown to increase risk for developmental delay and a variety of congenital anomalies in pediatric clinical populations ^27^. It is probable that some of the undiscovered rare alleles in SCZ are variants that confer protection but larger sample sizes are needed to determine this unequivocally. If true, our estimates of the excess CNV burden in cases may not fully account for the variation SCZ liability that is explained by rare CNVs.

Our results provide strong evidence that deletions in SCZ are enriched within a highly connected network of synaptic proteins, consistent with previous studies ^2,6,10,28^. The large CNV dataset here allows a more detailed view of the synaptic network and highlights subsets of genes account for the excess deletion burden in SCZ, including synaptic cell adhesion and scaffolding proteins, glutamatergic ionotropic receptors and protein complexes such as the ARC complex and DGC. Modest CNV evidence implicating Dystrophin (DMD) and its binding partners is intriguing given that the involvement of certain components of the DGC have been postulated ^29,30^ and disputed ^31^ previously. Larger studies of CNV are needed to define a role for this and other synaptic sub-networks in SCZ.

This study represents a milestone. Large-scale collaborations in psychiatric genetics have greatly advanced discovery through genome-wide association studies. Here we have extended this framework to rare CNVs. Our knowledge of the contribution from lower frequency variants gives us confidence that the application of this framework to large newly acquired datasets has the potential to further the discovery of loci and identification of the relevant genes and functional elements. The PGC CNV resource is now publicly available through a custom browser at http://pgc.tcag.ca/gb2/gbrowse/pgchg18/.

## Author Contributions

Management of the study, core analyses and content of the manuscript was the responsibility of the CNV Analysis Group chaired by J.S. and jointly supervised by S.W.S. and B.M.N. together with the Schizophrenia Working Group chaired by M.C.O’D. Core analyses were carried out by D.P.H., D.M., and C.R.M. Data Processing pipeline was implemented by C.R.M., B.T., W.W., D.G., M.G., A.S. and W.B. The A custom PGC CNV browser was developed by C.R.M, D.P.H., and B.T. Additional analyses and interpretations were contributed by W.W., D.A. and P.A.H. The individual studies or consortia contributing to the CNV meta-analysis were led by R.A.,O.A.A., D.H.R.B., A.D.B., E. Bramon, J.D.B., A.C., D.A.C., S.C., A.D., E. Domenici, H.E., T.E., P.V.G., M.G., H.G., C.M.H., N.I., A.V.J., E.G.J., K.S.K., G.K., J. Knight, T. Lencz, D.F.L., Q.S.L., J. Liu, A.K.M., S.A.M., A. McQuillin, J.L.M., P.B.M., B.J.M., M.M.N., M.C.O’D., R.A.O., M.J.O., A. Palotie, C.N.P., T.L.P., M.R., B.P.R., D.R., P.C.S, P. Sklar. D.St.C., P.F.S., D.R.W., J.R.W., J.T.R.W. and T.W. The remaining authors contributed to the recruitment, genotyping, or data processing for the contributing components of the meta-analysis. J.S., B.M.N, C.R.M, D.P.H., and D.M. drafted the manuscript, which was shaped by the management group. All other authors saw, had the opportunity to comment on, and approved the final draft.

## Acknowledgements

Core funding for the Psychiatric Genomics Consortium is from the US National Institute of Mental Health (U01 MH094421). We thank T. Lehner and Anjene Addington (NIMH). The work of the contributing groups was supported by numerous grants from governmental and charitable bodies as well as philanthropic donation. Details are provided in the Supplementary Notes. Membership of the Wellcome Trust Case Control Consortium and Psychosis Endophenotype International Consortium are provided in the Supplementary Notes.

### URLs

PGC CNV browser, http://pgc.tcag.ca/gb2/gbrowse/pgchg18.

---

**Extended data Table 1:** Previously reported CNV association. We assembled a list of 26 reported CNV associations to SCZ, where an odds ratio and *p*-value were available. At each CNV locus, we list the odds ratios and *p*-values from the largest sample collection available in the literature. Results from all CNV loci meta-analyzed in Rees et al. (2014), when available, were used. Throughout this article we refer to this entire list as “<u>previously reported</u>” loci. Reported *p*-values for nine loci shown in bold surpass the multiple testing threshold drawn from the current dataset using a Cochran-Mantel Haenszel test stratified by genotyping platform. Throughout this article we refer to these nine loci as “<u>previously implicated</u>”.

| Locus | CNV type | Gene or region name | Initial SCZ association reference (see legend) | Tested in Rees et al. 2014 | Reported *p*-value | SCZ CNV carrier % | Control CNV carrier % | Reported Odds Ratio |
| --- | --- | --- | --- | --- | --- | --- | --- | --- |
| **22q11.2** | deletion | multigenic | 1 | yes | **4.40E-40** | 0.29 | 0 | Inf |
| **16p11.2** | duplication | proximal duplication | 2 | yes | **2.90E-24** | 0.35 | 0.03 | 11.52 |
| **1q21.1** | deletion | multigenic | 3,4 | yes | **4.10E-13** | 0.17 | 0.021 | 8.35 |
| **2p16.3** | deletion | NRXN1 exons | 5,6 | yes | **1.30E-11** | 0.18 | 0.02 | 9.01 |
| **15q11.2** | deletion | multigenic | 3 | yes | **2.50E-10** | 0.59 | 0.28 | 2.15 |
| **3q29** | deletion | multigenic | 7,11 | yes | **1.50E-09** | 0.082 | 0.0014 | 57.65 |
| **15q13.2-13.3** | deletion | multigenic | 3,4 | yes | **5.60E-06** | 0.14 | 0.019 | 7.52 |
| **15q11.2-13.1** | duplication | AS/PWS | 8 | yes | **5.60E-06** | 0.083 | 0.0063 | 13.2 |
| **8q11.23** | duplication | RB1CC1 | 9 | no | **1.29E-05** | 0.106 | 0.014 | 8.58 |
| 16p13.11 | duplication | multigenic | 8 | yes | 5.70E-05 | 0.31 | 0.13 | 2.3 |
| 7q11.23 | duplication | Williams-Beuren | 10 | yes | 6.90E-05 | 0.066 | 0.0058 | 11.35 |
| 1q21.1 | duplication | multigenic | 11 | yes | 9.90E-05 | 0.13 | 0.037 | 3.45 |
| 16p13.2 | duplication | C16orf72/USP7 | 11 | no | 1.00E-04 | 0.254 | 0.0197 | 12.9 |
| 1p36.33 | duplication | multigenic | 12 | no | 5.00E-04 | 0.065 | 0.0075 | 8.66 |
| 22q11.2 | duplication | multigenic | 13 | no | 8.60E-04 | 0.014 | 0.085 | 0.17 |
| 17p12 | deletion | HNPP | 14 | yes | 1.20E-03 | 0.094 | 0.026 | 3.62 |
| 9q34.3 | duplication | intergenic | 15 | no | 1.40E-03 | 1.47 | 0.43 | 3.38 |
| 16p12.1 | deletion | multigenic | 12 | no | 1.60E-03 | 0.15 | 0.057 | 2.72 |
| 15q21.3 | duplication | CGNL1 | 12 | no | 1.90E-03 | 0.32 | 0.19 | 1.71 |
| 11q25 | deletion | GLB1L3/GLB1L2 | 11 | no | 3.00E-03 | 0.38 | 0.123 | 3 |
| 2q37.3 | duplication | AQP12A/KIF1A | 12 | no | 3.00E-03 | 0.34 | 0.24 | 1.43 |
| 17q12 | deletion | RCAD | 16 | yes | 0.0072 | 0.036 | 0.0054 | 6.64 |
| 9p24.2 | deletion | GLIS3 | 12 | no | 8.40E-03 | 0.033 | 0 | Inf |
| 9p24.2 | deletion | SLC1A1 | 12 | no | 9.80E-03 | 0.047 | 0.0075 | 6.19 |
| 16p11.2 | deletion | distal deletion | 17 | yes | 0.017 | 0.063 | 0.018 | 3.39 |
| 7q36.3 | duplication | WDR60/VIPR2 | 11,18 | yes | 0.27 | 0.11 | 0.069 | 1.54 |

**Extended Data Tables 2-4 are separate.xlsx sheets available upon request.**

**ED Table 2 datasets.xlsx** – datasets and sample sizes used in the current study

**ED Table 3 NeuroGeneSsets.xlsx** – Gene sets investigated in the current study

**ED Table 4 stepdown_withLegend.xlsx** – Unique contribution of significant gene sets in step-down regression models

**Extended Data Table 5:**
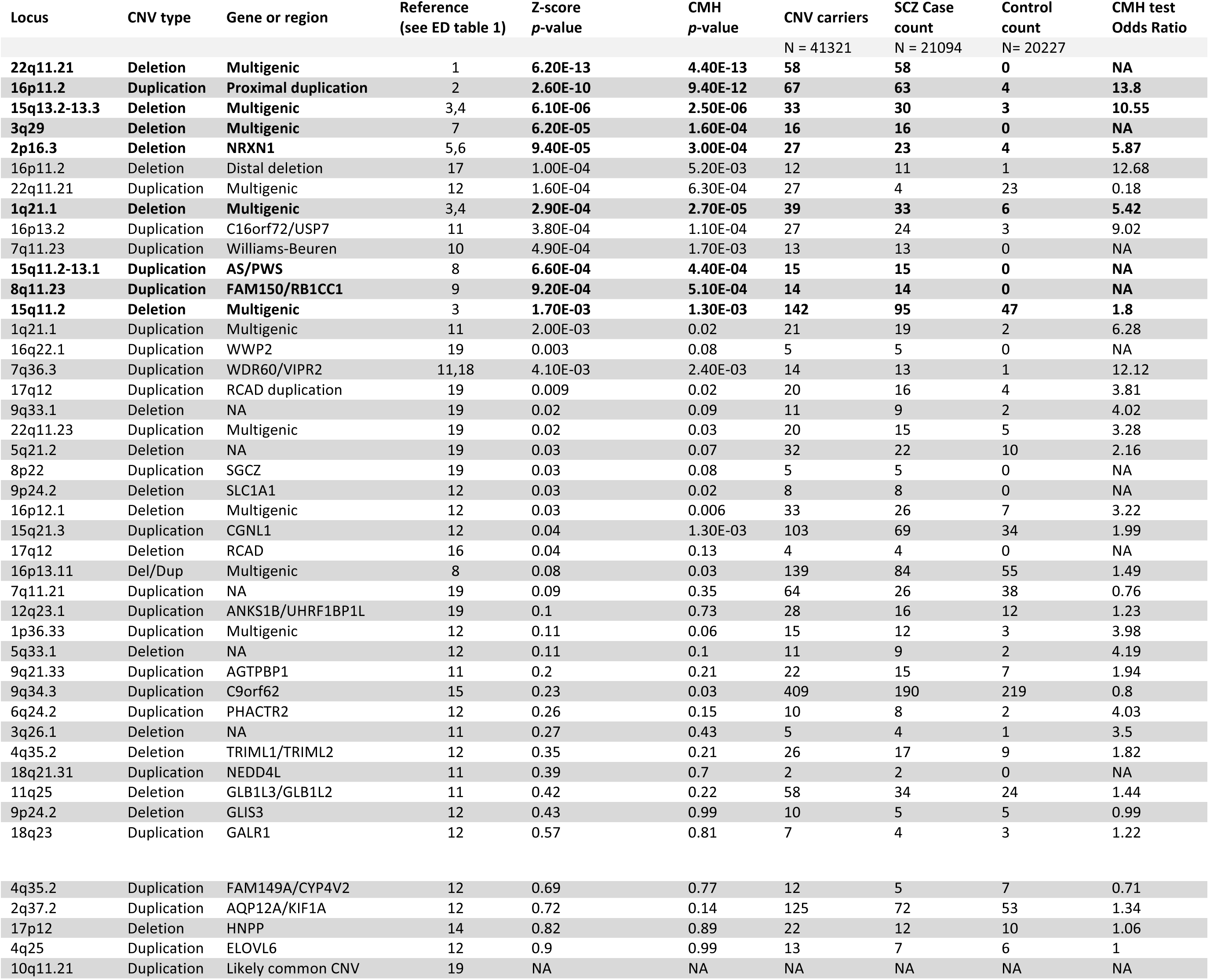
CNV probe-level results – Previously reported CNVs. Probe-level association results for all previously reported CNVs from genome-wide scans of SCZ. We report association results from the SCZ residual phenotype and from a CMH test stratified by genotyping platform. CNV loci in bold make up previously implicated loci, in which the most recent published *p*-value surpassed genome-wide correction.

| Locus | CNV type | Gene or region | Reference<br>(see ED table 1) | Z-score<br>p-value | CMH<br>p-value | CNV carriers<br>N = 41321 | SCZ Case<br>count<br>N = 21094 | Control<br>count<br>N = 20227 | CMH test<br>Odds Ratio |
| --- | --- | --- | --- | --- | --- | --- | --- | --- | --- |
| 22q11.21 | Deletion | Multigenic | 1 | 6.20E-13 | 4.40E-13 | 58 | 58 | 0 | NA |
| 16p11.2 | Duplication | Proximal duplication | 2 | 2.60E-10 | 9.40E-12 | 67 | 63 | 4 | 13.8 |
| 15q13.2-13.3 | Deletion | Multigenic | 3,4 | 6.10E-06 | 2.50E-06 | 33 | 30 | 3 | 10.55 |
| 3q29 | Deletion | Multigenic | 7 | 6.20E-05 | 1.60E-04 | 16 | 16 | 0 | NA |
| 2p16.3 | Deletion | NRXN1 | 5,6 | 9.40E-05 | 3.00E-04 | 27 | 23 | 4 | 5.87 |
| 16p11.2 | Deletion | Distal deletion | 17 | 1.00E-04 | 5.20E-03 | 12 | 11 | 1 | 12.68 |
| 22q11.21 | Duplication | Multigenic | 12 | 1.60E-04 | 6.30E-04 | 27 | 4 | 23 | 0.18 |
| 1q21.1 | Deletion | Multigenic | 3,4 | 2.90E-04 | 2.70E-05 | 39 | 33 | 6 | 5.42 |
| 16p13.2 | Duplication | C16orf72/USP7 | 11 | 3.80E-04 | 1.10E-04 | 27 | 24 | 3 | 9.02 |
| 7q11.23 | Duplication | Williams-Beuren | 10 | 4.90E-04 | 1.70E-03 | 13 | 13 | 0 | NA |
| 15q11.2-13.1 | Duplication | AS/PWS | 8 | 6.60E-04 | 4.40E-04 | 15 | 15 | 0 | NA |
| 8q11.23 | Duplication | FAM150/RB1CC1 | 9 | 9.20E-04 | 5.10E-04 | 14 | 14 | 0 | NA |
| 15q11.2 | Deletion | Multigenic | 3 | 1.70E-03 | 1.30E-03 | 142 | 95 | 47 | 1.8 |
| 1q21.1 | Duplication | Multigenic | 11 | 2.00E-03 | 0.02 | 21 | 19 | 2 | 6.28 |
| 16q22.1 | Duplication | WWP2 | 19 | 0.003 | 0.08 | 5 | 5 | 0 | NA |
| 7q36.3 | Duplication | WDR60/VIPR2 | 11,18 | 4.10E-03 | 2.40E-03 | 14 | 13 | 1 | 12.12 |
| 17q12 | Duplication | RCAD duplication | 19 | 0.009 | 0.02 | 20 | 16 | 4 | 3.81 |
| 9q33.1 | Deletion | NA | 19 | 0.02 | 0.09 | 11 | 9 | 2 | 4.02 |
| 22q11.23 | Duplication | Multigenic | 19 | 0.02 | 0.03 | 20 | 15 | 5 | 3.28 |
| 5q21.2 | Deletion | NA | 19 | 0.03 | 0.07 | 32 | 22 | 10 | 2.16 |
| 8p22 | Duplication | SGCZ | 19 | 0.03 | 0.08 | 5 | 5 | 0 | NA |
| 9p24.2 | Deletion | SLC1A1 | 12 | 0.03 | 0.02 | 8 | 8 | 0 | NA |
| 16p12.1 | Deletion | Multigenic | 12 | 0.03 | 0.006 | 33 | 26 | 7 | 3.22 |
| 15q21.3 | Duplication | CGNL1 | 12 | 0.04 | 1.30E-03 | 103 | 69 | 34 | 1.99 |
| 17q12 | Deletion | RCAD | 16 | 0.04 | 0.13 | 4 | 4 | 0 | NA |
| 16p13.11 | Del/Dup | Multigenic | 8 | 0.08 | 0.03 | 139 | 84 | 55 | 1.49 |
| 7q11.21 | Duplication | NA | 19 | 0.09 | 0.35 | 64 | 26 | 38 | 0.76 |
| 12q23.1 | Duplication | ANKS1B/UHRF1BP1L | 19 | 0.1 | 0.73 | 28 | 16 | 12 | 1.23 |
| 1p36.33 | Duplication | Multigenic | 12 | 0.11 | 0.06 | 15 | 12 | 3 | 3.98 |
| 5q33.1 | Deletion | NA | 12 | 0.11 | 0.1 | 11 | 9 | 2 | 4.19 |
| 9q21.33 | Duplication | AGTPBP1 | 11 | 0.2 | 0.21 | 22 | 15 | 7 | 1.94 |
| 9q34.3 | Duplication | C9orf62 | 15 | 0.23 | 0.03 | 409 | 190 | 219 | 0.8 |
| 6q24.2 | Duplication | PHACTR2 | 12 | 0.26 | 0.15 | 10 | 8 | 2 | 4.03 |
| 3q26.1 | Deletion | NA | 11 | 0.27 | 0.43 | 5 | 4 | 1 | 3.5 |
| 4q35.2 | Deletion | TRIML1/TRIML2 | 12 | 0.35 | 0.21 | 26 | 17 | 9 | 1.82 |
| 18q21.31 | Duplication | NEDD4L | 11 | 0.39 | 0.7 | 2 | 2 | 0 | NA |
| 11q25 | Deletion | GLB1L3/GLB1L2 | 11 | 0.42 | 0.22 | 58 | 34 | 24 | 1.44 |
| 9p24.2 | Deletion | GLIS3 | 12 | 0.43 | 0.99 | 10 | 5 | 5 | 0.99 |
| 18q23 | Duplication | GALR1 | 12 | 0.57 | 0.81 | 7 | 4 | 3 | 1.22 |
| 4q35.2 | Duplication | FAM149A/CYP4V2 | 12 | 0.69 | 0.77 | 12 | 5 | 7 | 0.71 |
| 2q37.2 | Duplication | AQP12A/KIF1A | 12 | 0.72 | 0.14 | 125 | 72 | 53 | 1.34 |
| 17p12 | Deletion | HNPP | 14 | 0.82 | 0.89 | 22 | 12 | 10 | 1.06 |
| 4q25 | Duplication | ELOVL6 | 12 | 0.9 | 0.99 | 13 | 7 | 6 | 1 |
| 10q11.21 | Duplication | Likely common CNV | 19 | NA | NA | NA | NA | NA | NA |

**Extended Data Figure 1:**
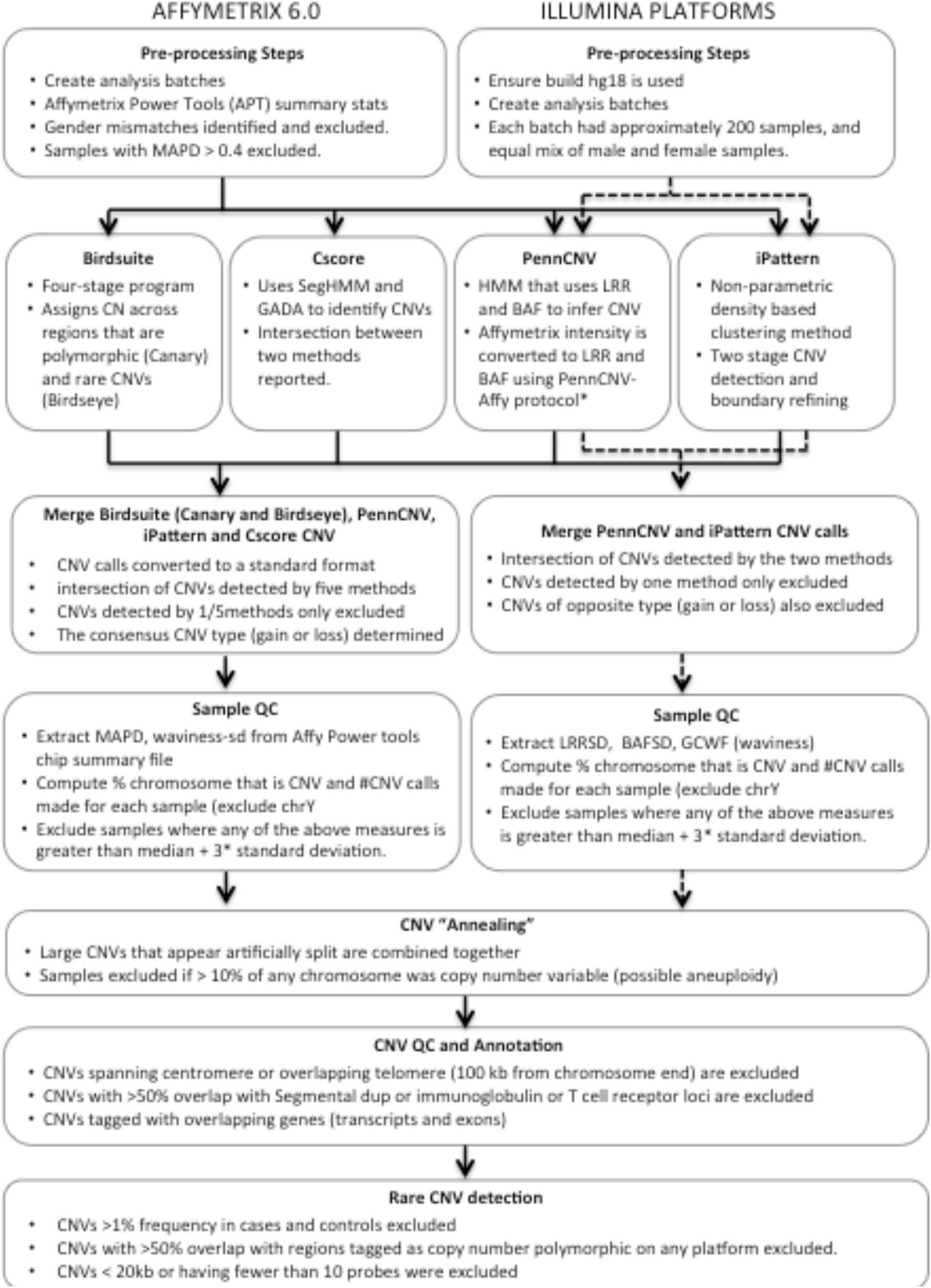
CNV pipeline workflow

**Extended Data Figure 2:**
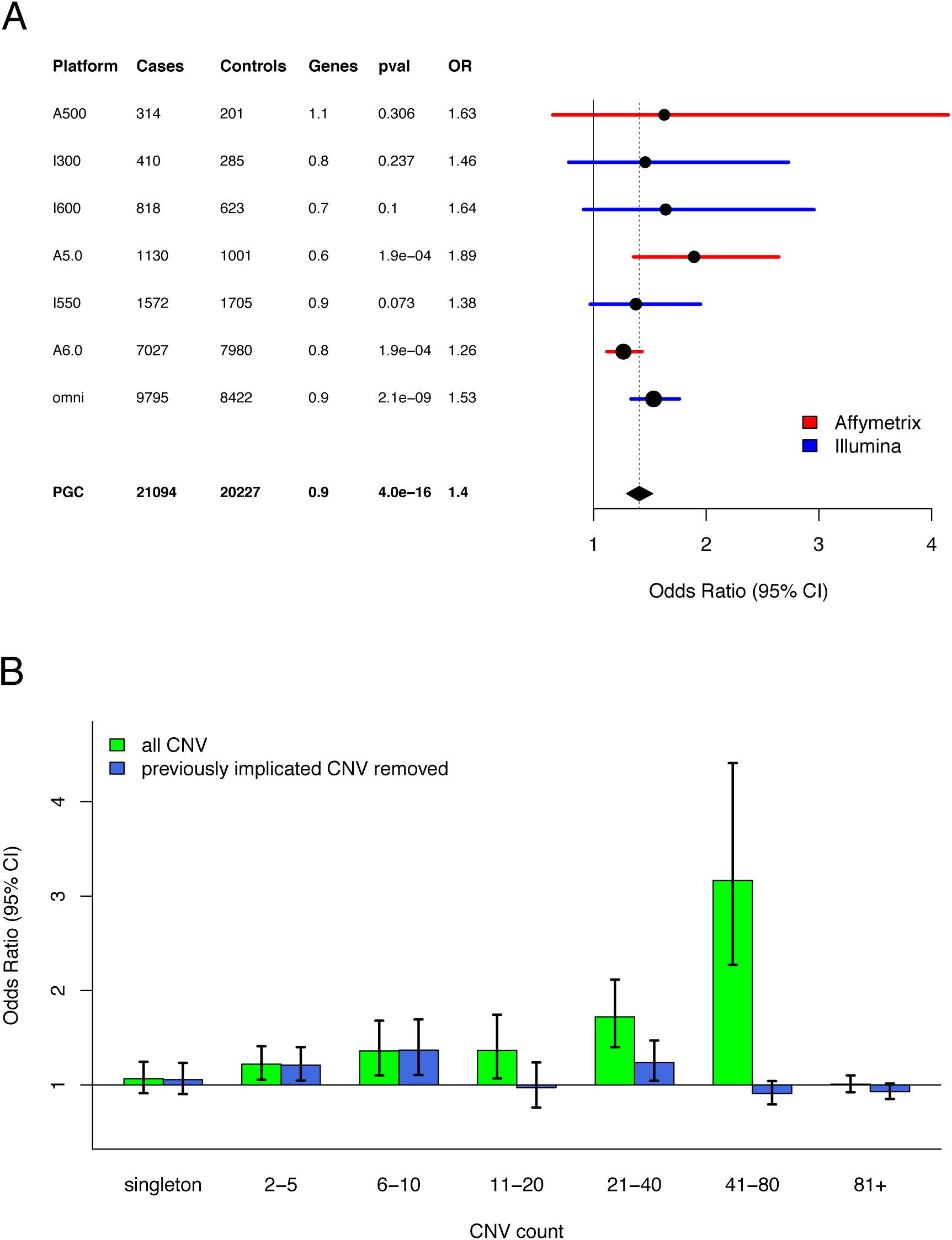
CNV burden for losses: Fig 3A: Forest plot of CNV burden (genes affected) partitioned by genotyping platform. Fig 3B: CNV burden partitioned by CNV frequency.

**Extended Data Figure 3:**
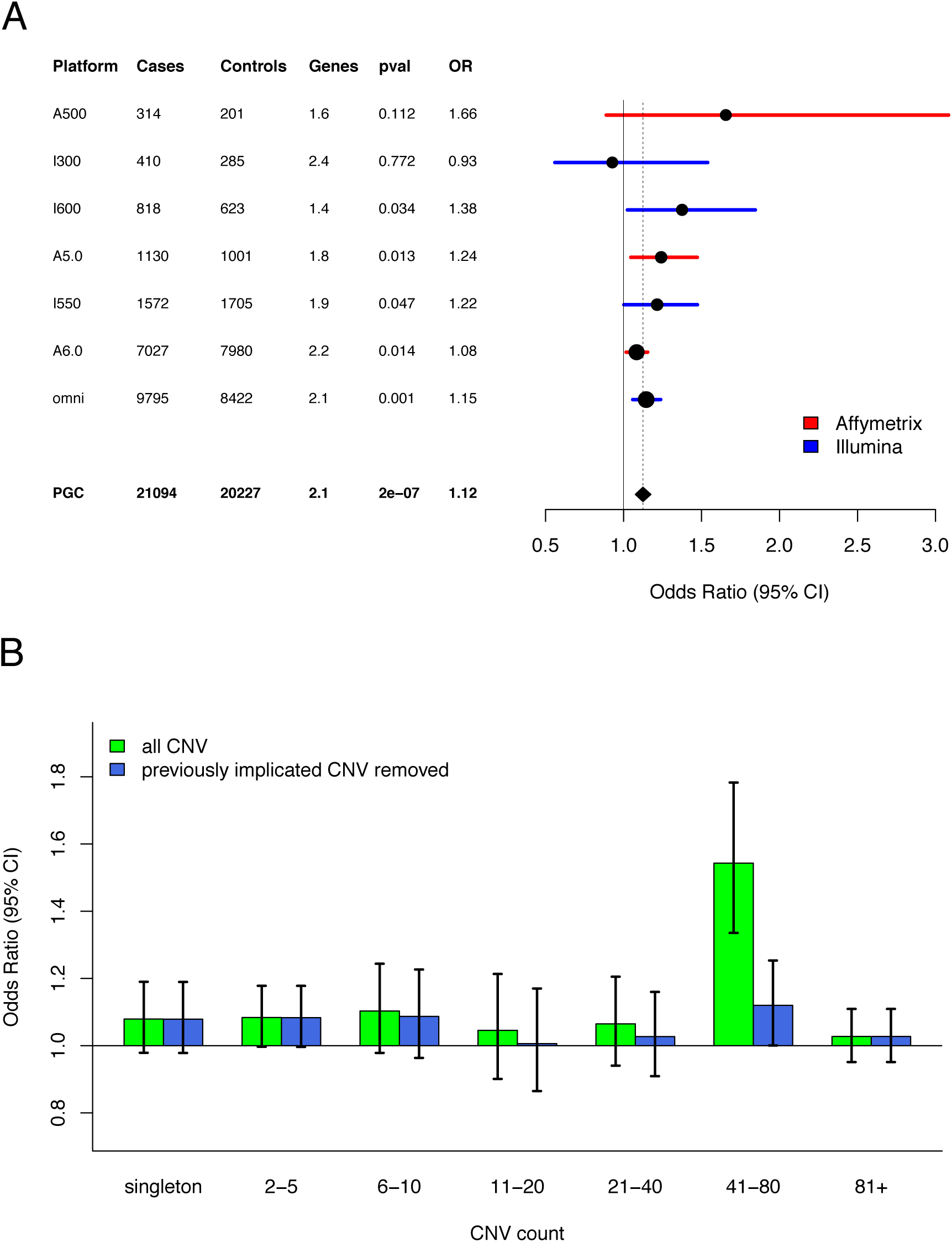
CNV burden for gains: Fig 4A: Forest plot of CNV burden (genes affected) partitioned by genotyping platform. Fig 4B: CNV burden partitioned by CNV frequency.

**Extended Data Figure 4:**
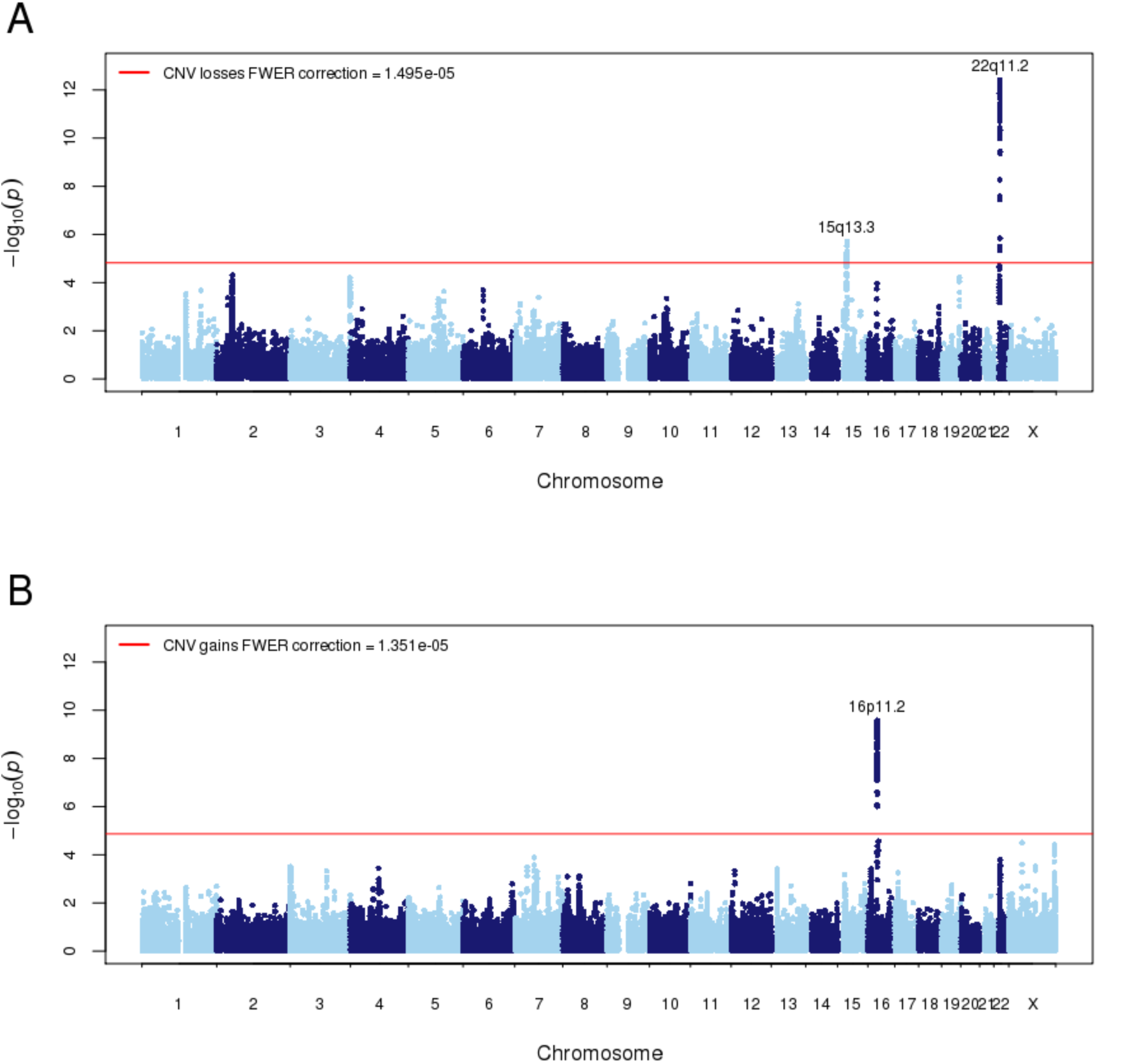
CNV probe level Manhattan plot: Manhattan plot of probe-level association results from the SCZ residual phenotype. Fig 5A: CNV losses Fig 5B: CNV gains. Genome-wide correction was determined using the family-wise error rate (FWER) drawn from permutation.

**Extended Data Figure 5:**
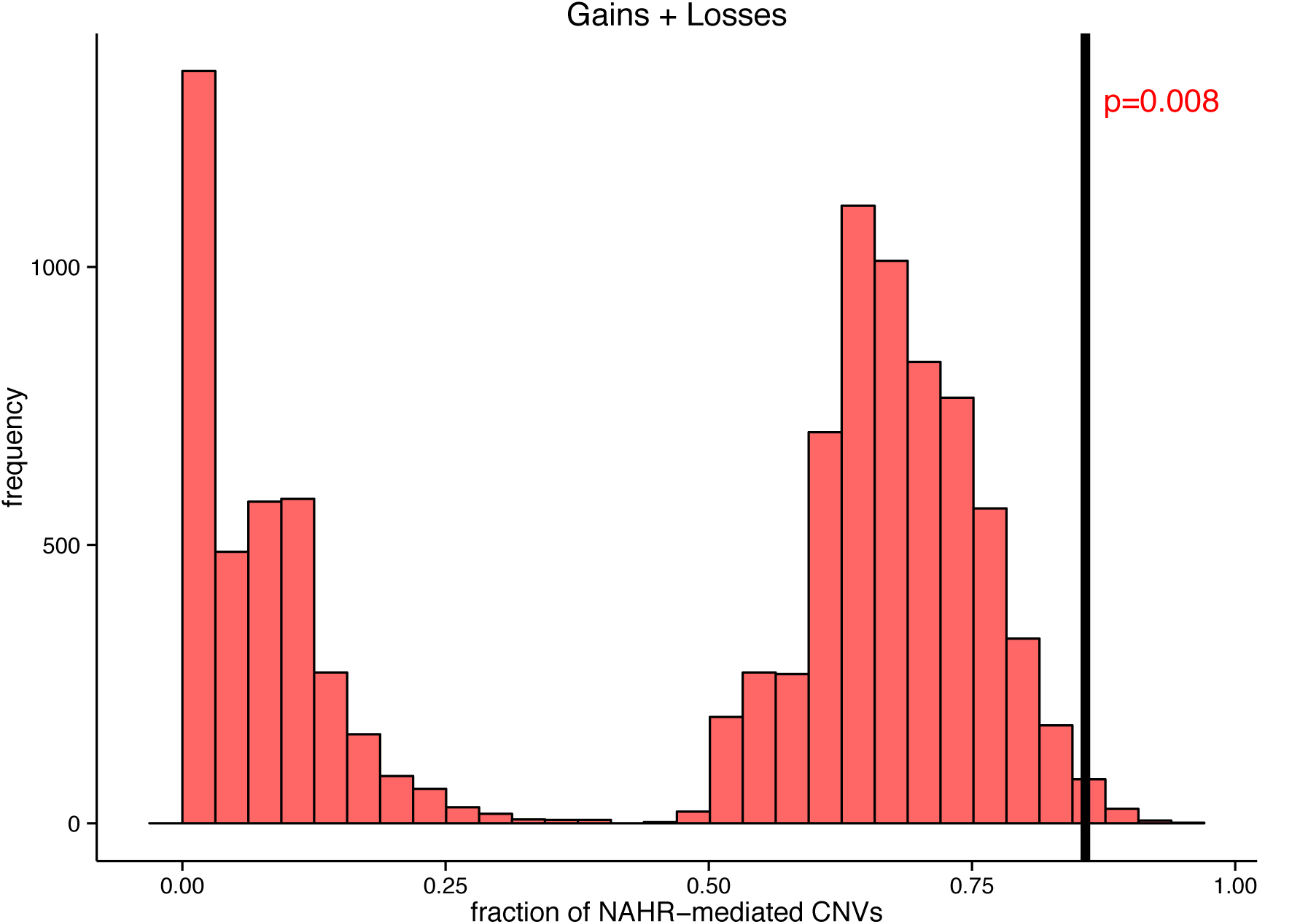
Permutation of NAHR-mediated CNVs: Permutation results from drawing frequency-match CNV loci and testing for fraction of NAHR-mediated CNVs. To test for the enrichment of NAHR-mediated loci in our suggestive results from the gene-based test, each permutation selected an equivalent number of independent CNV loci and tested the faction of NAHR-mediated CNVs.

**Extended Data Figure 6:**
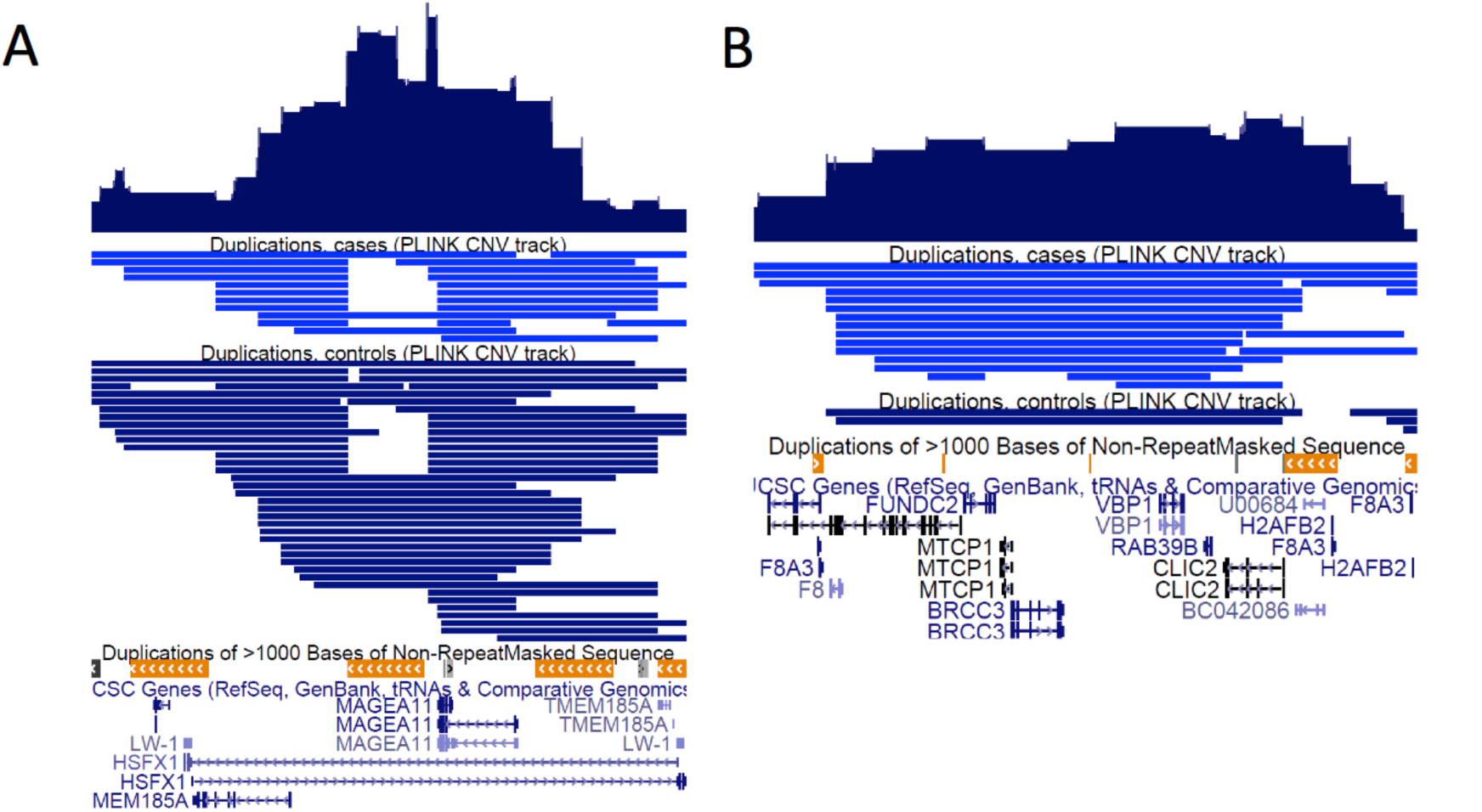
Xq28 CNV hotspot: Fig 6A: Protective CNV gain association peak around the MAGEA11 and TMEM185A gene, both within an intron of the HSFX1 gene. Fig 6B: Risk CNV gain association peak at the distal end of Xq28 overlapping ten genes.

**Extended Data Figure 7:**
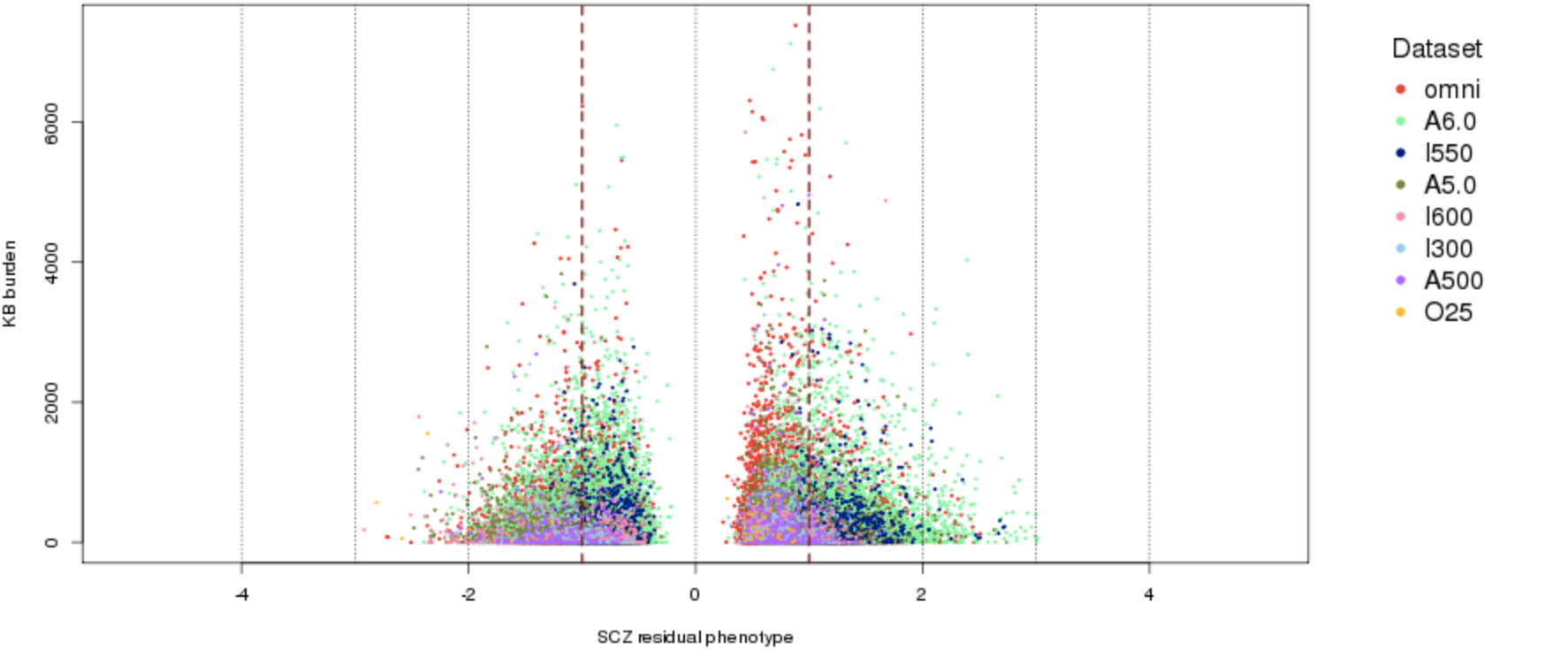
SCZ phenotype residual distribution: X-axis: Distribution of phenotype residual values after regressing case/control status on selected covariates. Plotted against overall CNV Kb burden (Y-axis) to visually inspect if individuals with large residuals have an excess of CNV burden, which can lead to higher false positive associations. SCZ cases have positive residual values and controls negative residual values.

**Extended Data Figure 8:**
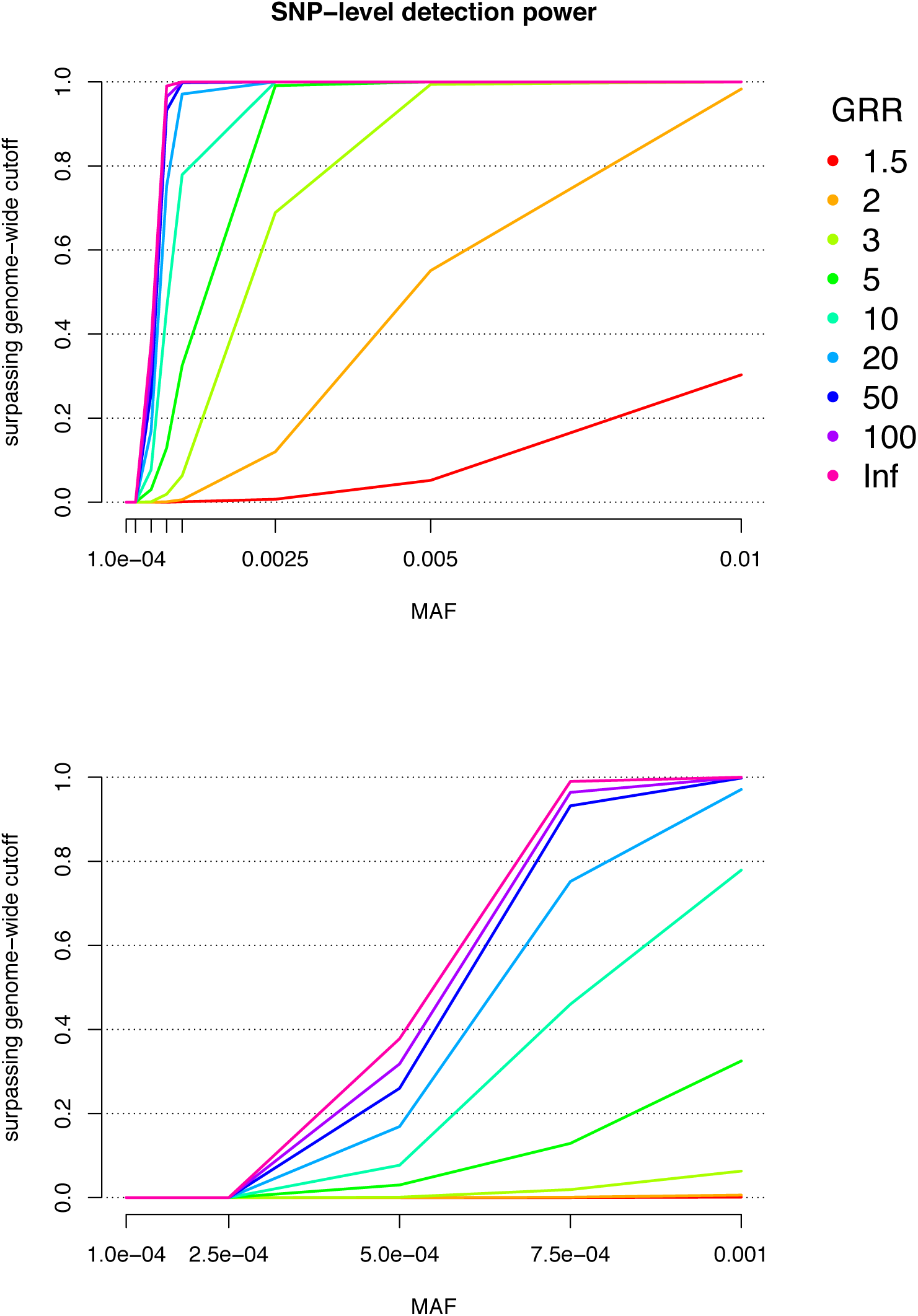
Detection power for CNV losses: Power is the proportion of simulated causal CNV loci detected (e.g. surpassing genome-wide FWER correction) using probe-level association. Each graph plots power across various MAF (x-axis) and genotype relative risk (GRR: colored lines). Simulations use the sample size and FWER cutoff from the current study.

## Methods

### Overview

We assembled a CNV analysis group with members from Broad Institute, Children’s Hospital of Philadelphia, University of Chicago, University of California San Diego, University of Michigan, University of North Carolina, Colorado University Boulder, and University of Toronto/SickKids Hospital. Our aim was to leverage the extensive expertise of the group to develop a fully automated centralized pipeline for consistent and systematic calling of CNVs for both Affymetrix and Illumina platforms. An overview of the analysis pipeline is shown in **Extended Data Figure 1**. After an initial data formatting step we constructed batches of samples for processing using four different methods, PennCNV, iPattern, C-score (GADA and HMMSeg) and Birdsuite for Affymetrix 6.0. For Affymetrix 5.0 data we used Birdsuite and PennCNV, for Affymetrix 500 we used PennCNV and C-score, and for all Illumina arrays we used PennCNV and iPattern. We then constructed a consensus CNV call dataset by merging data at the sample level and further filtered calls to make a final dataset **Extended data table 2**. Prior to any filtering, we processed raw genotype calls for a total of **57,577** individuals, including **28,684** SCZ cases and **28,893** controls.

### Study Sample

A complete list of datasets that were included in the current study can be found in **Extended Data Table 2**. A more detailed description of the original studies can be found in a previous publication^1^

### Copy Number Variant Analysis Pipeline Architecture and Sample Processing

All aspects of the CNV analysis pipeline were built on the Genetic Cluster Computer (GCC) in the Netherlands. PGC members sent external drives of raw data to the Netherlands for upload to the server as well as the corresponding sample metadata files.

#### Input Acceptance and Preprocessing

For Affymetrix we used the *.CEL files (all converted to the same format) as input, whereas for Illumina we required Genome or Beadstudio exported *.txt files with the following values: Sample ID, SNP Name, Chr, Position, Allele1 – Forward, Allele2 – Forward, X, Y, B Allele Freq and Log R Ratio. Samples were then partitioned into “batches’ to be run through each pipeline. For Affymetrix samples we created analysis batches based on the plate ID (if available) or genotyping date. Each batch had approximately 200 samples with an equal mix of male and female samples. Affymetrix Power Tools (APT - apt-copynumber-workflow) was then used to calculate summary statistics about chips analyzed. Gender mismatches identified and excluded as were experiments with MAPD > 0.4. For Illumina data, we first determined the genome build and converted to hg18 if necessary and created analysis batches based on the plate ID or genotyping date. Each batch had approximately 200 samples, and equal mix of male and female samples.

#### Composite Pipeline

The composite pipeline comprises CNV callers PennCNV ^2^, iPattern ^3^, Birdsuite ^4^ and C-Score ^5^ organized into component pipelines. We used all four callers for Affymetrix 6.0 data, PennCNV and C-Score for Affymetrix 500, Probe annotation files were preprocessed for each platform. Once the array design files and probe annotation files were pre-processed, each individual pipeline component pipeline was run in two steps: 1) processing the intensity data by the core pipeline process to produce CNV calls, 2) parsing the specific output format of the core pipeline and converting the calls to a standard form designed to capture confidence scores, copy number states and other information computed by each pipeline

### Merging of CNV data and Quality control filtering

#### Merging of CNV data

After standardization of outputs from each algorithm, CNV calls from each algorithm were merged at the sample level to increase specificity ^3^. For CNVs generated from Affymetrix 6.0 array, we took the intersection of the four outputs (Birdsuite, iPattern, C-Score, PennCNV) at the sample level to create a consensus CNV. For the Affymetrix 500, Affymetrix 5.0, and Illumina platforms, CNV merging was performed by taking the intersection of the calls made by the two algorithms (PennCNV and C-Score for Affymetrix 500, Birdsuite and PennCNV for Affymetrix 5.0, and iPattern and PennCNV for Illumina) at the sample level. CNV calls that were made by only one of the algorithm were excluded. Calls discordant for type of CNV (gain or loss) were also excluded.

#### Quality control filtering

Following merging we applied filtering criteria for removal of arrays with excessive probe variance or GC bias and removal of samples with mismatches in gender or ethnicity or chromosomal aneuploidies. For Affymetrix data, we extracted the MAPD and waviness-sd from the APT summary file. We also calculated the proportion of each chromosome (excluding chrY) tagged as copy number variable and computed the number of CNV calls made for each sample. We then retained experiments if each of these measures was within 3 SD of the median. For Illumina data, we extracted LRRSD, BAFSD, GCWF (waviness) from PennCNV log files. As with the Affymetrix data, we calculated the proportion of each chromosome (excluding chrY) tagged as copy number variable and computed the number of CNV calls made for each sample. We retained samples if each of the above measures was within 3 SD of the median. For both Illumina and Affymetrix datasets, large CNVs that appeared artificially split were combined together if one of the methods detected a CNV spanning the gap. However, samples where > 10% of the chromosome was copy number variable were excluded as possible aneuploidies. Further, we excluded CNVs that: 1) spanned the centromere or overlapped the telomere (100 kb from the ends of the chromosome); 2) had > 50% of its length overlapping a segmental duplication; 3) had >50% overlap with immunoglobulin or T cell receptor. The final filtered CNV dataset was annotated with Refseq genes (transcriptions and exons). After this stage of quality control (QC), we had a total of **52,511** individuals, with **27,034** SCZ cases and **25,448** controls.

#### Filtering for rare CNVs

To make our final dataset of rare CNVs for all subsequent analysis we universally filtered out variants that present at >= 1% (50% reciprocal overlap) frequency in cases and controls combined. CNVs that overlapped > 50% with regions tagged as copy number polymorphic on any other platform were also excluded. CNVs < 20kb or having fewer than 10 probes were also excluded.

### Post-CNV Calling QC

#### Overview

A number of steps were undertaken after CNV calling and initial filtering QC to minimize the impact of technical artifacts and potential confounds. In summary, we removed individuals not present in the PGC2 GWAS analysis ^1^, removed datasets with non-matching case or control samples that could not be reconciled using consensus platform probes, and removed any additional outliers with respect to overall CNV burden, CNV calling metrics, or SCZ phenotype residuals. All steps are described in more detail below.

#### Merging with GWAS cohort

By matching the unique sample identifiers, we retained only individuals that also passed QC filtering from the companion PGC GWAS study in Schizophrenia ^1^. This step filtered out samples with low-quality SNP genotyping, related individuals, and repeated samples across cohorts. An additional benefit of the PGC analytical framework is the ability to account for population stratification across cohorts using principal components derived from probe level analysis. After the post-CNV calling quality control steps described below, we re-calculated principal components using the Eigenstrat software package ^6^. Sample information and subsequent CNV and GWAS filtered sample sets are presented in **Extended data table 2**. In the process of matching to the GWAS-specific cohort, all individuals of non-European ancestry were removed from analysis (˜5.8% of the post-QC sample comprising three separate datasets). We also removed 42 samples that had discordant phenotype designations between the GWAS analysis and CNV genotype submission.

#### Individual dataset removal

Some datasets submitted to the PGC consisted of only case or control samples, affected trios, or recruited external samples as controls. This asymmetry in case-control ascertainment and genotyping can present serious biases for CNV analysis, as the sensitivity to detect CNV will vary considerably across genotyping platforms, as well as within dataset and genotyping batch. Unlike imputation protocols commonly used for SNP genotyping, there is no equivalent process to infer unmeasured probe intensity from nearby markers. We took a number of steps to identify and remove datasets that showed strong signs of case-control ascertainment or genotyping asymmetry:

1. Identify genotyping platforms where case-control ratio was not between 40-60%
2. Where possible, merge similar genotyping platforms using consensus probes prior to CNV-calling pipeline in order to improve case-control ratio.
3. Examine overall CNV burden and association peaks for spurious results
4. Remove datasets that remain problematic due to unusual CNV burden or multiple spurious CNV associations.

The genotyping platforms identified and processed are listed in **Extended data table 2**. We were able to combine the Illumina OmniExpress and Illumina OmniExpress plus Exome Chip platforms with success by removing probe content specific to the Exome chip platform. We removed the *caws* Affymetrix 500 datasets due to a number of strong CNV association peaks not seen in any other dataset. We also remove the *fii6* dataset due to a 2-fold CNV burden in cases relative to controls. In order to improve case-control balance, we had to remove the affected proband trio datasets (*boco, lacw*, and *lemu)* in the Illumina 610 platform, and the control-only *uclo* dataset in the Affymetrix 500 platform.

#### Individual sample removal

We re-analyzed CNV burden estimates in the reduced sample to flag any lingering outliers missed in the initial QC. We identified outliers for CNV count and Kb burden in the autosome (> 30 CNVs or 8 Mb, respectively) and in the X chromosome (> 10 CNVs or 5 Mb, respectively), removing an additional 15 individuals.

Genome-wide CNV intensity and quality measurements produced by CNV calling algorithms (i.e. “CNV metrics”) were examined for additional outliers and potential relationships with case-control status. Each CNV metric was re-examined across studies to assess if any additional outliers were present. Only three outliers were removed as their mean B allele (or minor allele) frequency deviated significantly from 0.5. Many CNV metrics are auto-correlated, as they measure similar patterns of variation in the probe intensity. Thus, we focused on the main intensity metrics - median absolute pairwise difference (MAPD) for projects genotyped on the Affymetrix 6.0 platform, and Log R Ratio standard deviation (LRRSD) in all other genotyping platforms. Among Affymetrix 6.0 datasets, MAPD did not differ between in cases and controls (t=1.14, *p* = 0.25). However, among non-Affymetrix 6.0 datasets, LRRSD showed significant differences between cases and controls (t=-35.3, *p* < 2e^−16^), with controls having a higher standardized mean LRRSD (0.227) than cases (-0.199). To control for any spurious associations driven by CNV calling quality, we included LRRSD (MAPD for Affymetrix 6.0 platforms) as a covariate in downstream analysis. CNV metrics were normalized with their genotyping platform prior to inclusion in the combined dataset.

### Regression of potential confounds on case-control ascertainment

The PGC cohorts are a combination of many datasets drawn from the US and Europe, and it is important to ensure that any bias in sample ascertainment does not drive spurious association to SCZ. In order to ensure the robustness of the analysis, we controlled for a number of covariates that could potential confound results. Burden and gene-set analyses included covariates in a logistic regression framework. Due to the number of tests run at probe level association, we employed a step-wise logistic regression approach to allow for the inclusion of covariates in our case-control association, which we term the *SCZ residual* phenotype.

Covariates include sex, genotyping platform, CNV metrics, and ancestry principal components derived from SNP genotypes on the same samples in a previous study^1^. We were unable to control for dataset or genotyping batch, as a subset of the contributing datasets are fully confounded with case/control status. CNV metric is normalized within genotyping platform prior to inclusion in the logistic model. Only principal components that showed a significant association to small CNV burden were used (small CNV being defined as autosomal CNV burden with CNV < 100 kb in size). Among the top 20 principal components, only the 1^st^, 2^nd^, 3^rd^, 4^th^, and 8^th^ principal component showed association with small CNV burden (with *p* < 0.01 used as the significance cutoff). To calculate the SCZ residual phenotype, we first fit a logistic regression model of covariates to affection status, and then extracted the Pearson residual values for use in a quantitative association design for downstream analyses. Residual phenotype values in cases are all above zero, and controls below zero, and are graphed against overall kb burden in **Extended data figure 7**. We removed three individuals with an SCZ residual phenotype greater than three (or negative three in controls). After the post-processing round of QC, we retained a dataset with a total of **41,321** individuals comprising **21,094** SCZ cases and **20,227** controls.

### Identifying previously implicated CNV loci in the literature

To delineate CNV burden effects coming from CNV loci that have previously been reported as putative SCZ risk factors from CNV in remainder of the genome, we flagged CNV loci with *p* < 0.01 that have either been reviewed ^7,8^ or otherwise reported ^8-10^ as potential SCZ risk factors in the literature. Previously reported loci meeting inclusion are listed in **Extended data table 1**. While a number of CNV loci have been reported in multiple studies, we sought the most recent reports that incorporated the largest sample sizes. To identify putatively associated CNV loci with SCZ from the full list, we applied the genome-wide *p*-value cutoff of **8e**^−5^, derived from the Cochran-Mantel-Haenzel (CMH) test in the current probe-level analysis as the *p*-value cutoff for inclusion as SCZ implicated CNV loci. While the CMH test is not the primary probe-level test in the current PGC analysis, it corresponds more closely to the tests used in published reports. In all, nine independent CNV loci from published reports surpass genome-wide correction. All published CNV loci, even those excluded as an SCZ implicated regions, are examined in the probe-level association analysis.

### CNV burden analysis

We analyzed the overall CNV burden in a variety of ways to discern which general properties of CNV are contributing to SCZ risk. Overall individual CNV burden was measured in 3 distinct ways – 1) Kb burden of CNVs, 2) Number of genes affected by CNVs, and 3) Number of CNVs. In particular, we only counted gene as affected when the CNV overlapped a coding exon. We also partitioned our analyses by CNV type, size, and frequency. CNV type is defined as copy number losses (or deletions), copy number gains (or duplications), and both copy number losses and gains. To assign a specific allele frequency to a CNV, we used the–cnv-freq-method2 command in PLINK, whereby the frequency is determined as the total number of CNV overlapping the target CNV segment by at least 50%. This method differs from other methods that assign CNV frequencies by genomic region, whereby a single CNV spanning multiple regions may be included in multiple frequency categories.

For **Figure 1**, and **Extended data figures 2** **and** **3**, we partitioned CNV burden by genotyping platform, and the abbreviations for each platform are expanded below:

A500: Affymetrix 500

I300: Illumina 300K

I600: Illumina 610K and Illumina 660W

A5.0: Affymetrix 5.0

A6.0: Affymetrix 6.0

omni: OmniExpress and OmniExpress plus Exome

Due to the small size of the Omni 2.5 array (28 cases and 10 controls), they were excluded from presentation in the figure, but are included in all burden analyses with the total PGC sample. Burden tests use a logistic regression framework with the inclusion of covariates detailed above. Using a logistic regression framework, we predicted SCZ status using CNV burden as an independent predictor variable, thus allowing us to get an accurate estimate of the unique contribution of CNV burden in a multiple regression framework. To gain insight into the proportion of CNV burden risk coming from loci outside of the previously implicated SCZ regions, we ran all burden analyses after removing CNV that overlapped previously implicated CNV boundaries by more than 10%.

### CNV probe level association

Genome-wide interrogation of CNV signals was tested at each respective CNV. Probe level tests were examined at the start, end, and single base position after the end of the called CNV. Three categories of CNV were tested: CNV deletions, CNV duplications, and deletions and duplications together. All analyses were run using PLINK software ^11^.

We ran probe level association using the SCZ residual phenotype as a quantitative variable, with significance determined through permutation of phenotype residual labels. An additional z-scoring correction, explained below, is used to control for any extreme values in the SCZ residual phenotype and efficiently estimate two-sided empirical *p*-values for highly significant loci. To ensure against the potential loss of power from the inclusion of covariates, we also ran a single degree of freedom Cochran-Mantel-Haenzel (CMH) test stratified by genotyping platform, with a 2 (CNV carrier status) x 2 (phenotype status) x N (genotyping platform) contingency matrix. While the CMH test does not account for more subtle biases that could drive false positive signals, it is robust to signals driven by a single platform and allows for each CNV carrier to be treated equally. Loci the surpassed genome-wide correction in either test was followed up for further evaluation.

#### Z-score recalibration of empirical testing

Probe level association *p*-values from the SCZ residual phenotype were initially obtained by performing one million permutations at each CNV position, whereby each permutation shuffles the SCZ residual phenotype among all samples, and retains the SCZ residual mean for CNV carriers and non-carriers. For extremely rare CNV, however, CNV carriers at the extreme ends of the SCZ residual phenotype can produce highly significant *p*-values. While we understand that such rare events are unable to surpass strict genome-wide correction, we wanted to retain all tests to help delineate the potential fine-scale architecture within a single region of association. To properly account for the increased variance when only a few individuals are tested, we applied an empirical Z-score correction to the CNV carrier mean. In order to get an empirical estimate of the variance for each test, we calculated the standard deviation of residual phenotype mean differences in CNV carriers and non-carriers from 5,000 permutations. Z-scores are calculated as the observed case-control mean difference divided by the empirical standard deviation, with corresponding *p*-values calculated from the standard normal distribution. Concordance of the initial empirical and z-score *p*-values are close to unity for association tests with six or more CNV, whereas Z-score *p*-values are more conservative among tests with less than six CNV. Furthermore, the Z-score method naturally provides an efficient manner to estimate highly significant empirical *p*-values that would involve hundreds of millions of permutations to achieve.

### Genome-wide correction for multiple tests

Beyond identifying significant CNV at the probe level, we also estimated the genome-wide testing space for rare CNV analysis. With the large PGC cohort being called through a consistent pipeline, we saw an opportunity to characterize the null expectation of segregating and recurrent *de novo* rare CNV in populations of European ancestry.

Accepted thresholds for significance among published risk CNV have been limited in scope, as accurate population estimates of rare CNV frequency and distribution across the genome require large representative samples.

Genome-wide significance thresholds were calculated using the 5% family-wise error rate from 5,000 permutations in both the SCZ residual phenotype and CMH test. Specifically, we selected the 95^th^ percentile of the minimum *p*-values obtained across permutations. Below are the genome-wide correction *p*-value thresholds determined in this manner:

*SCZ residual phenotype FWER correction:*

CNV losses and gains: 6.73e^−6^

CNV losses: 1.5e^−5^

CNV gains: 1.35e^−5^

*CMH test FWER correction:*

CNV losses and gains: 3.65e^−5^

CNV losses: 8.25e^−5^

CNV gains: 7.8e^−5^

This method differs slightly from those used in **Levinson et al.** ^9^ to estimate the multiple test correction for rare CNV, however their genome-wide correction of *p* = 1e^−5^ corresponds quite closely to the estimates observed using the SCZ residual phenotype. The observed family-wise correction serves as good approximation of the independent rare CNV signals found among European ancestry populations for array-based CNV capture, but as sample sizes increase, so too will the effective number of tests, necessitating further evaluation of the multiple testing burden.

### Gene-set burden enrichment analysis: gene-sets

Gene-sets with an a priori expectation of association to neuropsychiatric disorders were compiled based on gene annotations (Gene Ontology and curated pathway databases, downloaded June 2013) and published article materials (for details, see **Extended Data Table 3**). Gene-sets based on brain expression were compiled by processing the BrainSpan RNA-seq gene expression data-set (http://www.brainspan.org/static/download.html, downloaded Sept 2012). Four roughly equally sized gene-sets (about 4600 genes each) were derived to represent four expression tiers (very high, medium-to-high, medium-to-low, very low or absent); genes were selected if they passed a fixed expression threshold in at least 5/508 experimental data points (corresponding to different regions of donor brains, different donor ages corresponding to different developmental brain stages, and different donor sexes). Gene-sets based on mouse phenotypes were assembled by downloading MPO (Mammalian Phenotype Ontology) annotations from MGI (www.informatics.jax.org, downloaded August 2013), up-propagating annotations following ontology relations, and mapping to human orthologs using NCBI Homologene (www.ncbi.nlm.nih.gov/homologene); finally, top-level organ systems with fewer genes were aggregated while striving to preserve biological homogeneity, so to have roughly equal-sized sets (2,600-1,300 genes). For all gene-sets, gene identifiers in the primary source were mapped to Entrez-gene identifiers using the R/Bioconductor package *org.Hs.eg.db*.

### Gene-set burden enrichment analysis: pre-processing

Subjects were restricted to the ones with at least one rare CNV. For copy number gains and losses, we separately calculated the following subject-level totals: variant number, variant length and number of genes impacted; these covariates are then used to model global burden and correct gene-set burden to ensure it is specific (i.e. not a mere reflection of genome-wide burden with some stochastic deviation due to sampling). The subject-level total number of genes impacted was also calculated for each gene-set, again separately for gains and losses. Subjects were flagged if they carried at least one CNV matching a locus previously implicated in schizophrenia (see section “Identifying previously implicated CNV loci in the literature”); this was then used to analyzed gene-set burden for all subjects, or excluding subjects with an already implicated CNV.

### Gene-set burden enrichment analysis: statistical test

For each gene-set, we fit the following logistic regression model (as implemented by the R function *glm* of the *stats* package), where subjects are statistical sampling units: *y* ˜ *covariates + global + gene-set* Where:

- *y* is the dicotomic outcome variable (schizophrenia = 1, control = 0)
- *covariates* is the set of variables used as covariates also in the genome-wide burden and probe association analysis (sex, genotyping platform, CNV metric, and CNV associated principal components)
- global is the measure of global burden; for the results in the main text, we used the total gene number (abbreviated as *U* from universe gene-set count); we also calculated results for total length (abbreviated as *TL*) and variant number plus variant mean length (abbreviated as *CNML*)
- *gene-set* is the gene-set gene count

The gene-set burden enrichment was assessed by performing a chi-square deviance test (as implemented by the R function *anova.glm* of the *stats* package) comparing these two regression models:

*y* ˜ *covariates + global*

*y* ˜ *covariates + global + gene-set*

We reported the following statistics:

- coefficient beta estimate (abbreviated as *Coeff*)
- t-student distribution-based coefficient significance p-value (as implemented by the R function *summary.glm* of the *stats* package, abbreviated as *Pvalue_glm*)
- deviance test p-value (abbreviated as *Pvalue_dev*)
- gene-set size (i.e. number of genes is the gene-set, regardless of CNV data)
- BH-FDR (Benjamini-Hochberg False Discovery rate)
- percentage of schizophrenia and control subjects with at least 1 gene, 2 genes, etc … impacted by a CNV of the desired type (loss or gain) in the gene-set (abbreviated as *SZ_g1n, SZ_g2n, … CT_g1n*, …)

Please note that, by performing simple simulation analyses, we realized that *Pvalue_glm* can be extremely over-conservative in presence of very few gene-set counts different than 0, while *Pvalue_dev* tends to be slightly under-conservative. While the two p-values tend to agree well for gene-set analysis, *Pvalue_glm* is systematically over-conservative for gene analysis since smaller counts are typically available for single genes.

### Gene burden analysis: pre-processing

Subjects were restricted to the ones with at least one rare CNV. Only genes with at least a minimum number of subjects impacted by CNV were tested; this threshold was picked by comparing the BH-FDR to the permutation-based FDR and ensuring limited FDR inflation (permuted FDR < 1.65 * BH-FDR at BH-FDR threshold = 5%) while maximizing power. For gains the threshold was set to 12 counts, while for losses it was set to 8 counts.

### Gene burden analysis: statistical test

For each gene, we fit the following logistic regression model (as implemented by the R function *glm* of the *stats* package), where subjects are statistical sampling units: *y* ˜ *covariates + gene*

Where:

- *y* is the dichotomous outcome variable (schizophrenia = 1, control = 0)
- *covariates* is the set of variables used as covariates also in the genome-wide burden and probe association analysis (sex, genotyping platform, CNV metric, and CNV associated principal components)
- *gene* is the binary indicator for the subject having or not having a CNV of the desired type (loss or gain) mapped to the gene

The gene burden was assessed by performing a chi-square deviance test (as implemented by the R function *anova.glm* of the *stats* package) comparing these two regression models:

- *y* ˜ *covariates*
- *y* ˜ *covariates + gene*

### Gene burden analysis: multiple test correction

Multiple test correction was performed for loci rather than for genes, to avoid the strong correlation between test introduced by multi-genic CNVs; for the same reason, it is more useful to count false positives as loci rather than genes. We followed a greedy step-down procedure:

- start from gene with most significant deviance p-value G1, create locus L1
- remove from the gene list all genes that share at least 50% of their carrier subjects with G1, and add them to locus L1
- do the same for the next gene most significant gene in the list (thus creating a new locus L2), and proceed recursively until there is no gene left
- define locus p-value as the smallest deviance p-value of its genes

We computed permutation-based FDR by permuting subjects’ condition labels (schizophrenia, control), but not covariates (as those are expected to correlate to CNV distribution), 1,000 times. The FDR was then defined as the ratio between the average number of tests passing a given p-value threshold across the 1,000 permutations and the number of tests passing the same p-value threshold for real data. FDRs were also generated counting only the subset of genes with positive and negative regression coefficients (i.e. risk and presumed protective). The p-value threshold for permutation-based FDR calculation was picked by choosing the maximum nominal p-value corresponding to a given BH-FDR threshold (e.g. 5%). BH-FDR is supposed to be slightly inflated because (i) the deviance test p-value is slightly under-conservative in presence of very few gene indicators different than 0, (ii) we use the smallest gene p-value to define the locus p-value.

## PGC Schizophrenia CNV analysis – Supplementary Information

**<u>Supplementary Results:</u>**

- **CNV burden between sexes**
- **Probe level power analysis**
- **Gene-based network analysis**
- **Follow up of significant CNV loci**
- **Proportion of variance in SCZ explained by top CNV loci**
- **NAHR enrichment in significant novel gene loci**

**Consortium Membership**

**Acknowledgements**

**Supplementary Results**

## CNV burden between sexes

Following recent evidence that ostensibly healthy females carry an increased burden of rare CNVs ^1^, we examined whether this increased female burden existed in the current PGC dataset. We used a logistic regression model predicting sex using CNV burden and controlling for study covariates, as well as the Wilcoxon rank-sum test comparing male to female CNV burden ^1^. Focusing on the significant findings in the previous paper, we examined the burden in autosomal CNV count and genes affected among PGC controls (9856 males and 10371 females). We do see an elevated CNV count in control females (1.90 autosomal CNV rate) to males (1.87 autosomal CNV rate), however this difference is not significant in either the regression model (OR = 1.004, *p* = 0.66) or the Wilcoxon rank-sum test (*p* = 0.1). We do, however, observe a marginally significant enrichment when focusing on CNV loss count, where control females (0.99 autosomal CNV loss rate) show a higher burden than control males (0.94 autosomal CNV loss rate; logistic regression OR = 1.03, *p* = 0.05; Wilcoxon rank-sum test *p* = **3e**^−3^). No single genotyping platform seemed to drive the enrichment in females (data not shown), and we don’t observe any difference in CNV count when looking at CNV gains (logistic regression OR = 0.98, *p* = 0.18; Wilcoxon rank-sum test *p* = 0.56). Finally, no significant differences between sexes were found using either test when examining the number of genes affected, or when we include SCZ cases and controls (all *p* > .05).

## Probe level power analysis

By restricting analysis to rare CNV in the population (MAF < 0.01), many loci do not have enough CNV to surpass genome-wide correction for multiple testing, prompting pathway and gene level analyses to achieve sufficient statistical power. To use a specific example, the 3q29 deletion is fully penetrant in the current sample, with 16 SCZ carriers and 0 controls (MAF = **3.8e**^−4^) at the peak of association. Assuming no platform bias, this leads to an uncorrected chi-square *p*-value of **8.9e**^−5^, and a permuted *p*-value of **6.2e**^−5^ testing association using SCZ phenotype residuals. Neither *p*-value, however, surpasses their respective genome-wide significance cutoff for CNV deletion. While permutation methods used to generate genome-wide cutoffs accurately reflect the testing space among observed CNVs (very rare CNVs have little to no contribution to the family-wise error rate), we wanted to estimate the proportion of CNV detectable at the probe level. Under our current analytical design and sample size, we calculated the power to detect associated CNV across various MAFs and effect sizes and determine the proportion of association tests capable of surpassing genome-wide correction.

We simulated CNVs within our dataset (21094 cases and 20227 controls) and regressed them using the same association design with SCZ residual phenotypes. We simulated various effect sizes by randomly sampling cases and controls at different probabilities as CNV carriers, and rounded to the nearest CNV count to reflect the MAF of each CNV in the sample. For each combination of effect size and MAF, we ran 1000 simulations, retrieving the *t*-test *p*-value of CNV carriers from the SCZ residual phenotype. Simulated *p*-values behaved in much the same way as the Z-score correction on permutated *p*-values used in the primary test (data not shown). In **Extended data figure 8**, we show the proportion of simulations for CNV losses surpassing genome-wide correction at each MAF and effect size parameter (gains perform similarly).

We define statistical power as the proportion of simulations surpassing genome-wide significance. For a fully penetrant risk CNV, we require a MAF of ˜6e^−4^ (or about 25 CNV) to achieve 80% detection power. For CNV with a genotype relative risk (GRR) of 10, we require a MAF of 1e^−3^ (or at least 41 CNV) to achieve 80% detection power. Looking across the landscape of CNVs tested, on the whole about 10% of deletion or duplication CNV breakpoints reach a frequency greater than 1e^−3^ in the sample. On the other extreme, a CNV with MAF of .005 (or at least 206 CNV) and a GRR of 2 will only be detected 58% of the time.

## Gene-based network analysis

To identify a gene network enriched in schizophrenia risk genes, we queried GeneMANIA ^2^ using the 17 genes with deletion gene-test Benjamini-Hochberg FDR <= 25% and member of the “GO synaptic” or “ARC complex” sets. We thus created a synaptic protein interaction network of 136 genes, with the most densely connected network core corresponding to post-synaptic density organizers (DLGs, DLGAPs, SHANKs) and ionotropic glutamate receptors (GRIAs, GRIDs, GRINs). NRXN1 is connected to the network core via adhesion partners (NLGN1-3) and CASK. We tested this schizophrenia gene network, and found significant enrichment in genes with evidence of de novo coding variants in sequencing studies of schizophrenia trios ^3^ (for frameshift, stop-gain and splice-site: Fisher’s Exact Test p-value 0.0023; missense and amino acid insertion/deletion: Fisher’s Exact Test p-value 0.0004); in addition, we found a greater enrichment for this network, compared to the larger set composed of all “GO synaptic” and “ARC complex” genes. No significant enrichment was found for de novo variants identified in controls.

## Follow up of significant CNV loci

Both gene and probe level association follow a uniform testing framework across the genome, however risk loci may exhibit a more nuanced CNV architecture across the entirety of the association peak. All associated loci with FDR < .05 in the gene based test were followed up for further testing, along with a small number of candidate loci showing suggestive association in the probe-level association. We visually inspected each association peak and determined the bp coordinates that encapsulate the associated region and determine which CNV segment inclusion, be it covering exons or overlapping a minimum percentage of the total region, most appropriately reflect the association signal. To comprehensively examine the robustness and source of association, we also ran additional tests controlling for individual dataset, splitting by sex, and examining a dosage model, whereby copy number is measured with one copy for deletion, two copies for no CNV, and three copies for duplication. We also examined significant CNV loci in an unfiltered CNV call set, using CNVs called prior to the removal of common CNVs (MAF > 1%) and CNV overlapping segmental duplications.

We further evaluated the associated regions by determining the concordance of calls within the call set with those determined by unsupervised clustering. Call set CNVs were defined as CNVs with at least a 50% overlap with regions in **Table 1**. We restricted this analysis to 26,959 samples across six cohorts (14,419 Affymetrix 6.0, 12,540 Illumina platforms; 1.1:1 case:control ratio). Features for clustering included the median logR ratio (mLRR) and the median logR ratio of the chromosome for which a locus resides in, controlling for large chromosomal abnormalities. We implemented Density-Based Spatial Clustering of Applications (DBSCAN) found in the python scikit-learn library (http://scikit-learn.org) because of high sensitivity to detect outliers in clusters. For each novel region and within each cohort, genotypes were assigned to every sample based on the DBSCAN defined cluster. The cluster with the highest number of samples was designated as reference and assumed to have a copy number of two. Other clusters were flagged as gain or loss based on the average regional mLRR and its relation to the reference regional mLRR. We removed clusters with average chromosomal mLRR outside 3 SD from the reference. CNVs were considered concordant if they were flagged non-reference by DBSCAN and present in the 41k call set, matching on CNV type. We applied a locus based call set concordance filter of >=70%; one region, NPY4R, failed to meet this requirement with a concordance of 0.1%. In addition, both proximal and distal loci of ZNF600 were removed due to batch effects, which we defined as a significant deviation from a Poisson distribution of call set calls per plate. Regions that passed both concordance and batch effect filters are reported in **Table 1**.

## Proportion of variance in SCZ explained by top CNV loci

To measure the proportion of variance explained on the liability scale of SCZ, we estimated the overall heritability of liability (or logRR genetic variance) explained by the eight CNV loci surpassing genome-wide significance. All eight loci were collapsed into a single signal. Two SCZ affected individuals were found to carry two CNVs in these loci, and their contribution was only counted once. In sum, we observed 298 SCZ patients with a CNV in these regions (1.4% of the total SCZ affected sample), and 29 controls (0.1%; CMH stratified OR = 10.1). To estimate the variance in SCZ liability explained by loci surpassing genome-wide correction, we calculated the heritability of liability using the INDI-V online tool (cnsgenomics.com/software) described in ^4^ using an overall disease risk of 1% and a sibling recurrence risk of 8.8 ^5^.

## NAHR enrichment in significant novel gene loci

To test if novel significant loci (FDR<0.05; **Table 1**) were enriched for NAHR events, we performed a permutation test (n=10,000) simulating the null distribution of NAHR-mediated CNVs for a set of random loci. Each simulation randomly selected nine loci taken from CNVs overlapping at least 50% to genes in the gene-set burden analysis. These nine random loci were matched according to CNV call frequency to the nine novel significant loci in Table 1. We then created windows for each start and end position for every overlapping CNV to a random locus. Start positions were expanded −50kb and +5kb, and end positions were expanded −5kb and +50kb. We flagged CNVs as NAHR-mediated when both start and end expanded windows overlapped to 1kb segmental duplications obtained from the hg18 build of the UCSC table browser (https://genome.ucsc.edu/cgi-bin/hgTables). Every iteration reported the fraction of NAHR-mediated CNVs; that is the ratio of CNVs flagged as NAHR to the total number of overlapping CNVs. We found and enrichment of NAHR mediated CNVs in significant novel loci when compared to the null distribution (86% NAHR-mediated, 6 fold enrichment, p=0.008).

## Consortium Membership

### Wellcome Trust Case-Control Consortium 2

#### Management Committee

Peter Donnelly 180,217, Ines Barroso 218, Jenefer M Blackwell 219,220, Elvira Bramon 196, Matthew A Brown 221, Juan P Casas 222,223, Aiden Corvin 5, Panos Deloukas 218, Audrey Duncanson 224, Janusz Jankowski 225, Hugh S Markus 226, Christopher G Mathew 227, Colin N A Palmer 228, Robert Plomin 9, Anna Rautanen 180, Stephen J Sawcer 229, Richard C Trembath 227, Ananth C Viswanathan 230,231, Nicholas W Wood 232.

#### Data and Analysis Group

Chris C A Spencer 180, Gavin Band 180, Céline Bellenguez 180, Peter Donnelly 180,217, Colin Freeman 180, Eleni Giannoulatou 180, Garrett Hellenthal 180, RichardPearson 180, Matti Pirinen 180, Amy Strange 180, Zhan Su 180, Damjan Vukcevic 180.

#### DNA, Genotyping, Data QC, and Informatics

Cordelia Langford 218, Ines Barroso 218, Hannah Blackburn 218, Suzannah J Bumpstead 218, Panos Deloukas 218, Serge Dronov 218, Sarah Edkins 218, Matthew Gillman 218, Emma Gray 218, Rhian Gwilliam 218, Naomi Hammond 218, Sarah E Hunt218, Alagurevathi Jayakumar 218, Jennifer Liddle 218, Owen T McCann 218, Simon C Potter 218, Radhi Ravindrarajah 218, Michelle Ricketts 218, Avazeh Tashakkori-Ghanbaria 218, Matthew Waller 218, Paul Weston 218, Pamela Whittaker 218, Sara Widaa 218.

#### Publications Committee

Christopher G Mathew 227, Jenefer M Blackwell 219,220, Matthew A Brown 221, Aiden Corvin 5, Mark I McCarthy 233, Chris C A Spencer 180.

### Psychosis Endophenotype International Consortium

Maria J Arranz 156,234, Steven Bakker 101, Stephan Bender 235,236, Elvira Bramon 156,237,238, David A Collier 8,9, Benedicto Crespo-Facorro 239,240, Jeremy Hall 134, Conrad Iyegbe 156, Assen V Jablensky 241, René S Kahn 101, Luba Kalaydjieva 102,242, Stephen Lawrie 134, Cathryn M Lewis 156, Kuang Lin 156, Don H Linszen 243, Ignacio Mata 239,240, Andrew M McIntosh 134, Robin M Murray 142, Roel A Ophoff 80, Jim Van Os 143,156, John Powell 156, Dan Rujescu 81,83, Muriel Walshe 156, Matthias Weisbrod 236, Durk Wiersma 244.217

Department of Statistics, University of Oxford, Oxford, UK. 218 Wellcome Trust Sanger Institute, Wellcome Trust Genome Campus, Hinxton, Cambridge, UK. 219 Cambridge Institute for Medical Research, University of Cambridge School of Clinical Medicine, Cambridge, UK. 220 Telethon Institute for Child Health Research, Centre for Child Health Research, University of Western Australia, Subiaco, Western Australia, Australia. 221 Diamantina Institute of Cancer, Immunology and Metabolic Medicine, Princess Alexandra Hospital, University of Queensland, Brisbane, Queensland, Australia. 222 Department of Epidemiology and Population Health, London School of Hygiene and Tropical Medicine, London, UK. 223 Department of Epidemiology and Public Health, University College London, London, UK. 224 Molecular and Physiological Sciences, The Wellcome Trust, London, UK. 225 Peninsula School of Medicine and Dentistry, Plymouth University, Plymouth, UK. 226 Clinical Neurosciences, St George's University of London, London, UK. 227 Department of Medical and Molecular Genetics, School of Medicine, King's College London, Guy's Hospital, London, UK. 228 Biomedical Research Centre, Ninewells Hospital and Medical School, Dundee, UK. 229 Department of Clinical Neurosciences, University of Cambridge, Addenbrooke's Hospital, Cambridge, UK. 230 Institute of Ophthalmology, University College London, London, UK. 231 National Institute for Health Research, Biomedical Research Centre at Moorfields Eye Hospital, National Health Service Foundation Trust, London, UK. 232 Department of Molecular Neuroscience, Institute of Neurology, London, UK. 233 OxfordCentre for Diabetes, Endocrinology and Metabolism, Churchill Hospital, Oxford, UK. 234 Fundació de Docència i Recerca Mútua de Terrassa, Universitat de Barcelona, Spain. 235 Child and Adolescent Psychiatry, University of Technology Dresden, Dresden, Germany. 236 Section for Experimental Psychopathology, General Psychiatry, Heidelberg, Germany. 237 Institute of Cognitive Neuroscience, University College London, London, UK. 238 Mental Health Sciences Unit, University College London, London, UK. 239 Centro Investigación Biomédica en Red Salud Mental, Madrid, Spain. 240 University Hospital Marqués de Valdecilla, Instituto de Formación e Investigación Marqués de Valdecilla, University of Cantabria, Santander, Spain. 241 Centre for Clinical Research in Neuropsychiatry, The University of Western Australia, Perth, Western Australia, Australia. 242 Western Australian Institute for Medical Research, The University of Western Australia, Perth, Western Australia, Australia. 243 Department of Psychiatry, Academic Medical Center, University of Amsterdam, Amsterdam, The Netherlands. 244 Department of Psychiatry, University Medical Center Groningen, University of Groningen, The Netherlands.

### Acknowledgements

Data Processing and Statistical analyses were carried out on the Genetic Cluster Computer (http://www.geneticcluster.org) hosted by SURFsara and financially supported by the Netherlands Scientific Organization (NWO 480-05-003) along with a supplement from the Dutch Brain Foundation and the VU University Amsterdam. The GRAS data collection was supported by the Max Planck Society, the Max-Planck-Förderstiftung, and the DFG Center for Nanoscale Microscopy & Molecular Physiology of the Brain (CNMPB), Göttingen, Germany. The Boston CIDAR subject and data collection was supported by the National Institute of Mental Health (1P50MH080272, RWM; U01MH081928, LJS; 1R01MH092380, TLP) and the Massachusetts General Hospital Executive Committee on Research (TLP). ISC – Portugal: CNP and MTP are or have been supported by grants from the NIMH (MH085548, MH085542, MH071681, MH061884, MH58693, and MH52618) and the NCRR (RR026075). CNP, MTP, and AHF are or have been supported by grants from the Department of Veterans Affairs Merit Review Program. The Danish Aarhus study was supported by grants from The Lundbeck Foundation, The Danish Strategic Research Council, Aarhus University, and The Stanley Research Foundation. Work in Cardiff was supported by MRC Centre (G0800509) and MRC Programme (G0801418) Grants, the European Community's Seventh Framework Programme (HEALTH-F2-2010-241909 (Project EU-GEI)), the European Union Seventh Framework Programme (FP7/2007-2013) under grant agreement n° 279227, a fellowship to JW from the MRC/Welsh Assembly Government and the Margaret Temple Award from the British Medical Association. We thank Novartis for their input in obtaining CLOZUK samples, and staff at The Doctor's Laboratory (Lisa Levett/ Andrew Levett) for help with sample acquisition and data linkage and in Cardiff (Kiran Mantripragada/Lucinda Hopkins) for sample management. CLOZUK and some other samples were genotyped at the Broad Institute (which has a separate acknowledgment) or by the WTCCC and WTCCC2 (WT (083948/Z/07/Z). We acknowledge use of the British 1958 Birth Cohort DNA (MRC: G0000934) and the Wellcome Trust (068545/Z/0/ and 076113/C/04/Z), the UK Blood Services Common Controls (UKBS-CC collection), funded by the WT (076113/C/04/Z) and by NIHR programme grant to NHSBT (RP-PG-0310-1002). Virginia Commonwealth University: BPR and KSK thank all the faculty of the Virginia Institute for Psychiatric and Behavioral Genetics for invaluable insights and discussions over many years. BSM, SAB, BTW, BW, KSK and BPR were supported by National Institute of Mental Health grant R01 MH083094 to BPR. Sample collection was supported by previous funding of National Institute of Mental Health grant R01 MH041953 to KSK and BPR. Genotyping was supported by National Institute of Mental Health grant R01 MH083094 to BPR, National Institute of Mental Health grant R01 MH068881 to BPR and Wellcome Trust Case Control Consortium 2 grant. We thank Novartis for their input in obtaining CLOZUK samples, and staff at The Doctor's Laboratory (Lisa Levett/ Andrew Levett) for help with sample acquisition and data linkage and in Cardiff (Kiran Mantripragada/Lucinda Hopkins) for sample management. Our work was supported by: Medical Research Council (MRC) Centre (G0800509; G0801418), the European Community's Seventh Framework Programme (HEALTH-F2-2010-241909 (Project EU-GEI)), the European Union Seventh Framework Programme (FP7/2007-2013) under grant agreement n° 279227, a fellowship to JW from the MRC/Welsh Assembly Government and the Margaret Temple Award from the British Medical Association. CLOZUK and some other samples were genotyped at the Broad Institute (which has a separate acknowledgment) or by the WTCCC and WTCCC2 (WT (083948/Z/07/Z). We acknowledge use of the British 1958 Birth Cohort DNA (MRC: G0000934) and the Wellcome Trust (068545/Z/0/ and 076113/C/04/Z), the UK Blood Services Common Controls (UKBS-CC collection), funded by the WT (076113/C/04/Z) and by NIHR programme grant to NHSBT (RP-PG-0310-1002). The recruitment of families in Bulgaria was funded by the Janssen Research Foundation, Beerse, Belgium. We are grateful to the study volunteers for participating in the Janssen research studies and to the clinicians and support staff for enabling patient recruitment and blood sample collection. Informed consent was obtained from all participants or their parents or guardians. We thank the staff in the Neuroscience Biomarkers Genomic Lab led by Reyna Favis at Janssen for sample processing and the staff at Illumina for genotyping Janssen DNA samples. We also thank Anthony Santos, Nicole Bottrel, Monique-Andree Franc, William Cafferty of Janssen Research & Development) for operational support. Funding from the Netherlands Organization for Health Research and Development (ZonMw), within the Mental Health program (to GROUP consortium for collecting patients and clinical data). High-Density Genome-Wide Association Study Of Schizophrenia In Large Dutch Sample (R01 MH078075 NIH/National Institute Of Mental Health PI: Roel A. Ophoff). The Danish Council for Strategic Research (Journ.nr. 09-067048); The Danish National Advanced Technology Foundation (Journ.nr. 001-2009-2); The Lundbeck Foundation (Journ.nr. R24-A3243); EU 7th Framework Programme (PsychGene; Grant agreement nr. 218251); EU 7th Framework Programme (PsychDPC; Grant agreement nr. 286213). The Wellcome Trust supported this study as part of the Wellcome Trust Case Control Consortium 2 project. E. Bramon holds a MRC New Investigator Award and a MRC Centenary Award. The TOP Study was supported by the Research Council of Norway (#213837, # 217776, # 223273), South-East Norway Health Authority (# 2013-123) and K.G. Jebsen Foundation. This work was supported by the Donald and Barbara Zucker Foundation, the North Shore – Long Island Jewish Health System Foundation, and grants from the Stanley Foundation (AKM), the National Alliance for Research on Schizophrenia and Depression (AKM), and the NIH (MH065580 to TL; MH001760 to AKM). SynSys, EU FP7-242167, Sigrid Juselius Foundation, The Academy of Finland, grant number: 251704, Sohlberg Foundation. The Swedish Research Council [grant numbers 2006-4472, 2009-5269, 2009-3413] and the County Councils of Västerbotten and Norrbotten, Sweden supported the collection of the scz_umeb_eur and scz_umes_eur samples. The Betula Study, from which the Umea controls were recruited, is supported by grants from the Swedish Research Council [grant numbers 345-2003-3883, 315-2004-6977] and the Bank of Sweden Tercentenary Foundation, the Swedish Council for Planning and Coordination of Research, the Swedish Council for Research in the Humanities and Social Sciences and the Swedish Council for Social Research. The GRAS (Göttingen Research Association for Schizophrenia) data collection has been supported by the Max Planck Society, the Max Planck Förderstiftung, and the DFG (CNMPB). We thank all GRAS patients for participating in the study, and all the many colleagues who have contributed over the past 10 years to the GRAS data collection. We acknowledge support from the North Shore – LIJ Health System Foundation and NIH grants RC2 MH089964 and R01 MH084098. We acknowledge support from NIMH K01 MH085812 (PI Keller) and NIMH R01 MH100141 (PI Keller). EGCUT work was supported by the Targeted Financing from the Estonian Ministry of Science and Education [SF0180142s08]; the US National Institute of Health [R01DK075787]; the Development Fund of the University of Tartu (grant SP1GVARENG); the European Regional Development Fund to the Centre of Excellence in Genomics (EXCEGEN; grant 3.2.0304.11-0312); and through FP7 grant 313010. Milan Macek was supported by CZ.2.16/3.1.00/24022OPPK, NT/13770-4and 00064203 FN Motol. For the scz_tcr1_asn dataset funding from the National Medical Research Council (Grant: NMRC/TCR/003/2008) and the Biomedical Research Council, A*STAR is acknowledged. Genotyping of the Swedish Hubin sample was performed by the SNP&SEQ Technology Platform in Uppsala, which is supported by Uppsala University, Uppsala University Hospital, Science for Life Laboratory - Uppsala and the Swedish Research Council (Contracts 80576801 and 70374401). The Swedish Hubin sample was supported by Swedish Research Council (IA, EGJ) and the regional agreement on medical training and clinical research between Stockholm County Council and the Karolinska Insititutet (EGJ). B.J.M., V.J.C., R.J.S., S.V.C., F.A.H., A.V.J., C.M.L., P.T.M., C.P., and U.S. were supported by the Australian Schizophrenia Research Bank, which is supported by an Enabling Grant from the National Health and Medical Research Council (Australia) [No. 386500], the Pratt Foundation, Ramsay Health Care, the Viertel Charitable Foundation and the Schizophrenia Research Institute and the NSW Department of Health. C.P. is supported by a Senior Principal Research Fellowship from the National Health and Medical Research Council (Australia). We acknowledge the help of: Johanna Badcock, Linda Bradbury, Jason Bridge, David Chandler, Janell Collins-Langworthy, Trish Collinson, Milan Dragovic, Cheryl Filippich, David Hawkes, Danielle Lowe, Kathryn McCabe, Tamara MacDonald, Barry Maher, Bharti Morar Marc Seal, Heather Smith, Melissa Tooney, Paul Tooney, and Melinda Ziino. The Danish Aarhus study was supported by grants from The Lundbeck Foundation, The Danish Strategic Research Council, Aarhus University, and The Stanley Research Foundation. The Perth sample collection was funded by Australian National Health and Medical Research Council project grants and the Australian Schizophrenia Research Bank. The Bonn/Mannheim sample was genotyped within a study that was supported by the German Federal Ministry of Education and Research (BMBF) through the Integrated Genome Research Network (IG) MooDS (Systematic Investigation of the Molecular Causes of Major Mood Disorders and Schizophrenia; grant 01GS08144 to M.M.N. and S.C., grant 01GS08147 to M.R.), under the auspices of the National Genome Research Network plus (NGFNplus), and through the Integrated Network IntegraMent (Integrated Understanding of Causes and Mechanisms in Mental Disorders), under the auspices of the e:Med Programme.(GSK control sample; Müller-Myhsok). This work has been funded by the Bavarian Ministry of Commerce and by the Federal Ministry of Education and Research in the framework of the National Genome Research Network, Förderkennzeichen 01GS0481 and the Bavarian Ministry of Commerce. M.M.N. is a member of the DFG-funded Excellence-Cluster ImmunoSensation. M.M.N. also received support from the Alfried Krupp von Bohlen und Halbach-Stiftung. M.R. was also supported by the 7th Framework Programme of the European Union (ADAMS project, HEALTH-F4-2009-242257; CRESTAR project, HEALTH-2011-1.1-2) grant 279227. Roche: Thanks are expressed to Olivia Spleiss for great support in genetic data generation, Daniel Umbricht and Delphine Lagarde for their valuable support in clinical and genetic data sharing, and Anirvan Ghosh for continuous encouragement. Authors also wish to thank all investigators and patients who participated in the Roche clinical studies. Jo Knight holds the Joanne Murphy Professor in Behavioural Science. We thank Maria Tampakeras for her work on the samples. The Stanley Center for Psychiatric Research at the Broad Institute acknowledges funding from the Stanley Medical Research Institute. Swedish schizophrenia study (PI CMM, PFS, PS, SM): We are deeply grateful for the participation of all subjects contributing to this research and to the collection team that worked to recruit them: E. Flordal-Thelander, A.-B. Holmgren, M. Hallin, M. Lundin, A.-K. Sundberg, C. Pettersson, R. Satgunanthan-Dawoud, S. Hassellund, M. Rådstrom, B. Ohlander, L. Nyrén and I. Kizling. Funding support for the Sweden Schizophrenia Study (PIs Hultman, Sullivan, and Sklar) was provided by the NIMH (R01 MH077139 to P.F.S. and R01 MH095034 to P.S.), the Stanley Center for Psychiatric Research, the Sylvan Herman Foundation, the Friedman Brain Institute at the Mount Sinai School of Medicine, the Karolinska Institutet, Karolinska University Hospital, the Swedish Research Council, the Swedish County Council, the Söderström Königska Foundation. We acknowledge use of DNA from The UK Blood Services collection of Common Controls (UKBS collection), funded by the Wellcome Trust grant 076113/CI04/Z, by the Juvenile Diabetes Research Foundation grant WT0618S8, and by the National Institute of Health Research of England. The collection was established as part of the Wellcome Trust Case-Control Consortium. We thank the study participants, and the research staff at the study sites. This study was supported by NIMH grant R01MH062276 (to DF Levinson, C Laurent, M Owen and D Wildenauer), grant R01MH068922 (to PV Gejman), grant R01MH068921 (to AE Pulver) and grant R01MH068881 (to B Riley). The authors are grateful to the many family members who participated in the studies that recruited these samples, to the many clinicians who assisted in their recruitment. In addition to the support acknowledged for the Multicenter Genetics Studies of Schizophrenia and Molecular Genetics of Schizophrenia studies, Dr. DF Levinson received additional support from the Walter E. Nichols, M.D., Professorship in the School of Medicine, the Eleanor Nichols Endowment, the Walter F. & Rachael L. Nichols Endowment and the William and Mary McIvor Endowment, Stanford University. This study was supported by NIH R01 grants (MH67257 to N.G.B., MH59588 to B.J.M., MH59571 to P.V.G., MH59565 to R.F., MH59587 to F.A., MH60870 to W.F.B., MH59566 to D.W.B., MH59586 to J.M.S., MH61675 to D.F.L., MH60879 to C.R.C., and MH81800 to P.V.G.), NIH U01 grants (MH46276 to C.R.C., MH46289 to C. Kaufmann, MH46318 to M.T. Tsuang, MH79469 to P.V.G., and MH79470 to D.F.L.), the Genetic Association Information Network (GAIN), and by The Paul Michael Donovan Charitable Foundation. Genotyping was carried out by the Center for Genotyping and Analysis at the Broad Institute of Harvard and MIT (S. Gabriel and D. B. Mirel), which is supported by grant U54 RR020278 from the National Center for Research Resources. Genotyping of half of the EA sample and almost all the AA sample was carried out with support from GAIN. The GAIN quality control team (G.R. Abecasis and J. Paschall) made important contributions to the project. We thank S. Purcell for assistance with PLINK. We (DRW, RS) thank the staff of the Lieber Institute and the Clinical Brain Disorders Branch of the IRP, NIMH for their assistance in data collection and management. We thank Ningping Feng and Bhaskar Kolachana for Illumina genotyping and for managing DNA stocks. The work was supported by the Lieber Institute and by direct NIMH IRP funding of the Weinberger Lab. Pfizer is very grateful to the study volunteers for participating in our research studies. We thank our numerous clinicians and support staff for enabling patient recruitment, blood sample collection, and biospecimen administration. Informed consent was obtained from all participants, their parents or guardians. Eli Lilly is grateful to the participants of clinical trials and research studies who gave consent for participation in this study. We are also grateful to Philip J Ebert and Jeffrey S Arnold for facilitating our participation in this project. We acknowledge the Irish contribution to the International Schizophrenia Consortium (ISC) study, the WTCCC2 SCZ study & WTCCC2 controls from the 1958BC and UKNBS, the Science Foundation Ireland (08/IN.1/B1916). We thank the Toronto Centre for Applied Genomics for technical and computational assistance and funding from the University of Toronto McLaughlin Centre and Genome Canada. S.W.S. holds the GlaxoSmithKline-CIHR Chair in Genome Sciences at the Hospital for Sick Children and University of Toronto. We acknowledge use of the Trinity Biobank sample from the Irish Blood Transfusion Service & the Trinity Centre for High Performance Computing. Funding for this study was provided by the Wellcome Trust Case Control Consortium 2 project (085475/B/08/Z and 085475/Z/08/Z), the Wellcome Trust (072894/Z/03/Z, 090532/Z/09/Z and 075491/Z/04/B), NIMH grants (MH 41953 and MH083094) and British 1958 Birth Cohort DNA collection funded by the Medical Research Council (grant G0000934) and the Wellcome Trust (grant 068545/Z/02) and of the UK National Blood Service controls funded by the Wellcome Trust. We acknowledge Hong Kong Research Grants Council project grants GRF 774707M, 777511M, 776412M and 776513M.

